# Genomic signatures of past megafrugivore-mediated dispersal in Malagasy palms

**DOI:** 10.1101/2023.01.20.524701

**Authors:** Laura Méndez, Christopher D. Barratt, Walter Durka, W. Daniel Kissling, Wolf L. Eiserhardt, William J. Baker, Vonona Randrianasolo, Renske E. Onstein

**Affiliations:** German Centre for Integrative Biodiversity Research (iDiv) Halle-Jena-Leipzig, Puschstrasse 4, 04103 Leipzig, Germany; Department of Community Ecology (BZF), Helmholtz Centre for Environmental Research - UFZ, Theodor-Lieser-Strasse 4, 06120 Halle, Germany; Institute for Biodiversity and Ecosystem Dynamics (IBED), University of Amsterdam, P.O. Box 94240, 1090 GE Amsterdam, The Netherlands; Royal Botanic Gardens, Kew, Richmond, Surrey, TW9 3AE, United Kingdom; Department of Biology, Aarhus University, Ny Munkegade 116, 8000 Aarhus C, Denmark; Kew Madagascar Conservation Centre, Ivandry, Madagascar; Naturalis Biodiversity Center, Darwinweg 2, 2333 CR Leiden, the Netherlands

**Keywords:** Arecaceae, ddRAD, genetic diversity, genetic differentiation, megafauna extinction, population connectivity, population genomics, seed dispersal

## Abstract

1. Seed dispersal is a key process in the generation and maintenance of genetic diversity and genetic differentiation of plant populations in tropical ecosystems. During the Last Quaternary, most seed-dispersing megafauna was lost globally, but whether this has caused dispersal limitation, loss of genetic diversity, and increased genetic differentiation between plant populations with large, ‘megafaunal’ fruits (i.e., > 4 cm - megafruits) remains unclear.
2. Here, we assessed whether megafrugivore extinctions in Madagascar (e.g., giant lemurs, elephant birds) have affected the genetic diversity and genetic differentiation of four animal-dispersed Malagasy palm (Arecaceae) species with large (*Borassus madagascariensis*), medium-sized (*Hyphaene coriacea, Bismarckia nobilis*), and small (*Chrysalidocarpus madagascariensis*) fruits. We integrated double-digest restriction-site-associated DNA sequencing (ddRAD) of 167 individuals from 25 populations with (past) distribution ranges for extinct and extant seed-dispersing animal species, climate and human impact data, and applied linear mixed-effects models to explore the drivers of variation in genetic diversity and genetic differentiation across Malagasy palm populations.
3. We detected higher genetic diversity in species with megafruits than in the species with small fruits, and genetic differentiation was lowest for the human-used medium-sized megafruit species. Furthermore, we found that a higher number of shared extinct megafrugivore species between palm population pairs was associated with less genetic differentiation, indicating higher gene flow, whereas no relationship with extant frugivores – that are not able to swallow and disperse the seeds – was found. Finally, genetic diversity decreased with road density, whereas genetic differentiation decreased with increasing human population density, but only for populations with megafruits.
4. Our results suggest that the legacy of megafrugivores regularly achieving long dispersal distances is still reflected in the genetic diversity and genetic differentiation of palms that were formerly dispersed by such large animals. Furthermore, high genetic diversity and low genetic differentiation were possibly maintained after the megafauna extinctions through human-mediated dispersal, long generation times, and long lifespans of these palms. Our study illustrates how integrating genetics with ecological data on species interactions, climate, and human impact, provides novel insights into the consequences of megafauna extinctions for plants with megafruits.

## Introduction

Understanding how genetic diversity and genetic differentiation are generated and maintained is crucial to predict species’ adaptive potential for evolutionary change. Genetic diversity and genetic differentiation depend on species-specific life-history traits, population dynamics, past climatic and demographic events, biogeography, and local and global environmental factors (Ellegren & Galtier, 2016). Specifically for plants, seed dispersal and pollination play key roles in population connectivity, allowing gene flow between non-adjacent populations, thereby controlling regional-scale genetic diversity and structure (Browne et al., 2018). Furthermore, seed dispersal allows plants to colonize new habitats through long-distance dispersal events. In tropical rainforests, more than 90% of woody plant species rely on frugivores (i.e., fruit-eating and seed-dispersing animals) for their seed dispersal (Jordano, 2000). For these plants, most long-distance dispersal events are provided by large-bodied frugivores (i.e., megafrugivores - subset of largest animal species having fruit as the main part of their diet in a given ecosystem; Moleón et al., 2020), which are able to ingest a wide range of fruit and seed sizes and move them across long distances given their large home ranges (Pires et al., 2018).

During the last 125,000 years, extinction rates of large-bodied animals have increased globally due to human impact (Smith et al., 2018). The extirpation of megafrugivores from ecosystems may have cascading effects on the seed dispersal and thus connectivity of plant populations, especially for plants carrying large fruits that cannot be dispersed by the remaining smaller-bodied frugivores in the ecosystem (Janzen & Martin, 1982). Frugivore extinctions may therefore directly affect plant genetic diversity and genetic differentiation (Giombini et al., 2017; Pérez-Méndez et al., 2016) and the capacity of vertebrate-dispersed plants to track climate change (Fricke et al., 2022). Though macroevolutionary inferences have indicated increased extinction rates across New World megafaunal-fruited palms in the Quaternary (Onstein et al., 2018), there remains no strong evidence of any plant species that went extinct in response to the extinction of its megafrugivore interaction partners. Instead, it has been suggested that several plant species that were adapted to seed dispersal by megafrugivores have persisted due to domestication by humans (Kistler et al., 2015), secondary seed dispersal by smaller-bodied frugivores (Blanco et al., 2019), or other, non-biotic forms of dispersal.

In Madagascar, megafauna abundances started to decrease around 1000 years ago, primarily due to increasing human impact (Crowley, 2010). Hunting and a transition to herding and farming, thereby changing landscapes and megafauna habitats, ultimately led to the extirpation of all megafauna (Godfrey et al., 2019; Li et al., 2020). Some of these extinct megafaunal species have been identified as fruit-eaters, such as giant lemurs (e.g. *Pachylemur* spp. and *Archaeolemur* spp.), elephant birds (e.g., *Aepyornis* spp. and *Vorombe* spp.) and giant tortoises (*Aldabrachelys* spp.) (Godfrey et al., 2004; Pedrono et al., 2013). Fossil pollen suggests that the extinction of endemic megafauna coincided with a gradual decline in abundance of trees relying on megafrugivores for seed dispersal (Domic et al., 2021). Moreover, Malagasy palm distributions still retain signals of past dispersal by extinct megafrugivore animals, mostly in the western part of the island (Méndez et al., 2022). However, the extent to which dispersal services provided by Madagascar’s past megafrugivores can still be detected in the genomes of extant plant populations remains unclear.

In addition to biotic interactions with frugivorous animals, there are other mechanisms such as isolation by distance (Wright, 1943), isolation by environment (Wang & Bradburd, 2014) and isolation by resistance (McRae, 2006) that influence the genetic diversity and genetic differentiation of plant populations (Jiang et al., 2019; Sexton et al., 2014; Siepielski et al., 2017). For example, current climate, specifically precipitation, had a small, negative relationship with genetic diversity across 384 plant species, leading to a weak latitudinal gradient in genetic diversity (i.e., lower genetic diversity towards the equator; De Kort et al., 2021). In Madagascar, rivers acted as barriers leading to high genetic structure in leafless vanilla orchids (*Vanilla* spp.), and both isolation by distance and isolation by environment were inferred to affect genetic differentiation between orchid populations (Andriamihaja et al., 2021). Current forest cover has also been shown to be important for driving genetic diversity and genetic structure of endemic plants in Madagascar (Salmona et al., 2022). Finally, besides frugivores and the abiotic environment, humans have been a major determinant of present-day genetic diversity and genetic structure of plants (Arredondo et al., 2018; Smith et al., 2020). For example, by moving plant seeds, humans can cause long-distance dispersal events (Wichmann et al., 2008), thereby increasing genetic diversity and decreasing genetic differentiation (Bullock et al., 2018). Moreover, fragmentation of ecosystems and human settlement can lead to local extinctions, decreased population sizes, loss of genetic diversity, and reduced movement of large vertebrates (e.g., mammals) with a subsequent loss of seed dispersal of plants with large vertebrate-dispersed fruits (Tucker et al., 2021).

Here, we assessed whether megafrugivore extinctions have affected the genetic diversity and genetic differentiation of megafaunal-fruited palm (Arecaceae) species in Madagascar, compared to a small-fruited palm. We focused on palms because several species have been identified as ‘anachronisms’ in Madagascar’s ecosystems due to their megafaunal fruit sizes (> 4 cm in length, hereafter referred to as megafruits; Guimarães et al., 2008) that seem maladapted to dispersal by the current frugivore pool (Albert-Daviaud et al., 2020). We selected species from the western part of the island, where megafrugivore animals were possibly most abundant in the past (Crowley, Godfrey, & Irwin, 2011). These savanna animal-dispersed palm species can be classified in three fruit size classes: large megafruits (30 cm in average lenght - *Borassus madagascariensis*), medium-sized megafruits (5.5 cm - *Hyphaene coriacea*, 4.4 cm - *Bismarckia nobilis*) and small fruits (< 4 cm; 1.3 cm - *Chrysalidocarpus madagascariensis* [previously *Dypsis madagascariensis*, Eiserhardt et al., 2022]) (Table 1). The relatively small fruits of *C. madagascariensis* can still be dispersed by extant frugivores (e.g., *Eulemur macaco*; Adany et al., 1994), whereas the megafruits of the other three species are too large to be swallowed and dispersed by any extant native frugivore on Madagascar (Perry & Hartstone-Rose, 2010). Although not dispersed by extant frugivores, both palms with medium-sized megafruits (*H. coriacea* and *B. nobilis*) are highly used by humans for house construction, basketry or food (Rakotoarinivo et al., 2020). Human dispersal may thus have contributed to the persistence and genetics of these species during the last thousand years. Finally, sexual system and pollination may also affect gene flow and thus population genetics. In terms of their sexual system, the three megafruit species included in the tribe Borasseae (*B. madagascariensis, H. coriacea* and *B. nobilis*) are dioecious, with male and female trees separated, while *C. madagascariensis* is monoecious (male and female flowers on the same tree). Little is known about the pollinators of these palms, but flowers are relatively small and inconspicuous, suggesting a wide range of insect pollinators (Henderson, 1986).

**Table 1.** Fruit size, number of populations and genetic diversity parameters averaged per species: Observed heterozygosity (Ho), expected heterozygosity (He) and nucleotide diversity (π). For more detailed information and values calculated per population, see Table S7 in supporting information.

| | Average fruit length (cm) | Average fruit width (cm) | Number of populations | $H_o$ | $H_e$ | $\pi$ |
| --- | --- | --- | --- | --- | --- | --- |
| *Borassus madagascariensis* | 30 | 12.75 | 3 | 0.245 | 0.230 | 0.250 |
| *Bismarckia nobilis* | 4.4 | 3.25 | 8 | 0.223 | 0.206 | 0.231 |
| *Hyphaene coriacea* | 5.5 | 5 | 7 | 0.244 | 0.232 | 0.259 |
| *Chrysalidocarpus madagascariensis* | 1.3 | 0.75 | 7 | 0.193 | 0.179 | 0.198 |

We hypothesize (H1) that the extinction of Madagascar’s megafrugivores has reduced seed dispersal, connectivity and thus gene flow between palm populations with megafruits (large and medium megafruits). We therefore expect lower genetic diversity and higher genetic differentiation in populations of species with megafruits (i.e., *B. madagascariensis, H. coriacea, B. nobilis)* than in populations of the species with small fruits (i.e., *C. madagascariensis*). However, if megafrugivores used to be important seed dispersers of the palm species with megafruits, historical long-distance dispersal events by megafrugivores may still be reflected in the genetic diversity and genetic differentiation of palm populations with megafruits. We therefore expect (H2) that populations that used to share (i.e., co-occurred with) the same megafrugivore species were more connected in the past, and thus genetically more similar (i.e., lower genetic differentiation) than populations that were less connected by past seed dispersal. Consequently, populations that used to co-occur with a higher megafrugivore species richness are also expected to harbour higher genetic diversity than populations that co-occurred with fewer megafrugivore species. In comparison, co-occurrence with extant frugivores is expected to not affect genetic diversity or genetic differentiation of palm populations with megafruits, because these large palm fruits supposedly do not interact with smaller-bodied extant frugivores. Furthermore, human use may additionally affect genetic diversity and genetic differentiation of palms. We therefore expect (H3) that higher road density and higher human population density have increased gene flow between populations of the two human-used medium-sized megafruit palms, and thus increased genetic diversity and decreased genetic differentiation between populations. Finally, we explored the influence of abiotic climatic factors (e.g., precipitation and temperature seasonality, proximity to rivers and forest cover), isolation by distance, and isolation by resistance on genetic diversity and genetic differentiation of Malagasy palm populations. For specific predictions on genetic diversity, genetic differentiation and genetic structure for each hypothesized scenario, see Figure 1.

**Figure 1.**
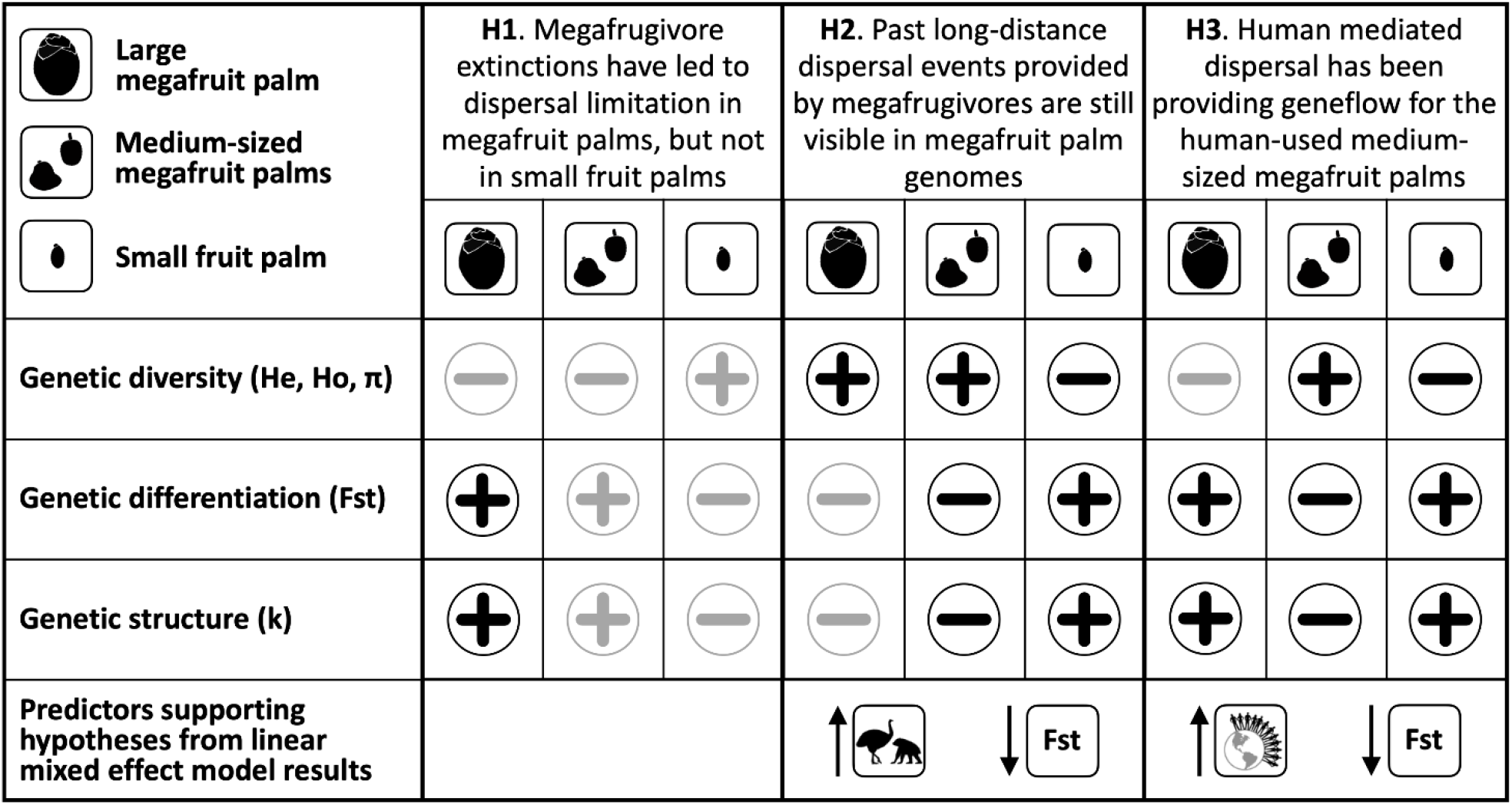
Overview of all main hypotheses and specific predictions for each fruit class on genetic diversity, genetic differentiation and genetic structure. A plus symbol indicates the expectation of a relatively (compared to other fruit sizes) high value for that genetic measurement, and a minus symbol indicates the expectation of a low value for that genetic measurement. When the plus and minus symbols appear in black, it means that the prediction was supported by our results; when the symbol appears in grey, it means that the prediction was not supported by our results. We also show the specific predictors from the linear mixed effect models results that supported each hypothesis, and the general trend it showed with population genetic differentiation (Fst). Symbol of frugivores refers to shared number of extinct megafrugivores between population pairs. Symbol of human refers to accumulative human population density between population pairs. See Figure 5 for more details on the results of the linear mixed effect models.

To test these hypotheses, we integrated palm population-level sampling of genomic regions with past inferences of megafrugivore distributions, current frugivore distributions, climate and human impact variables, species distribution modelling and phylogenetic reconstructions. We applied linear mixed-effects models to disentangle the frugivory-related, human, and abiotic effects on the genetics of Malagasy palms. We therefore provide novel insights in the consequences of megafrugivore extinctions for plants on Madagascar.

## Materials and methods

### Sample collection and library preparation

We sampled leaf tissue from 25 natural populations of four palm species that differ in fruit sizes (large megafruits: *B. madagascariensis*; medium-sized megafruits: *H. coriacea, B. nobilis*; small fruits: *C. madagascariensis*) throughout their distribution in the western part of Madagascar during July, August and September of 2019 (Figure 2, Table 1, Table S1). More details on sampling for each species are provided in Table 1 and Tale S1.

**Figure 2.**
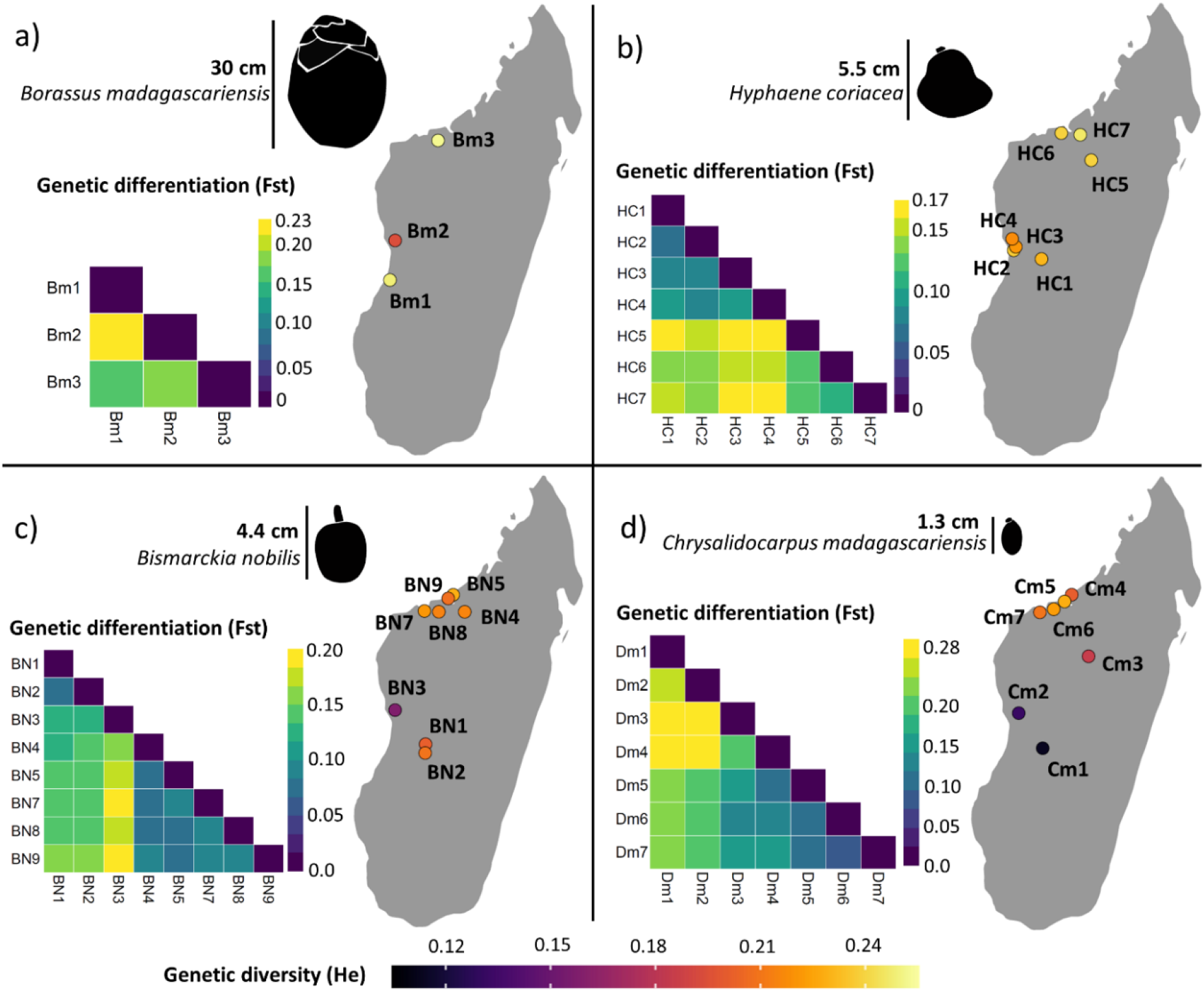
Map of Madagascar showing the different populations sampled for four palm species, and their genetic diversity and genetic differentiation. Brighter colours reflect higher genetic diversity (expected heterozygosity – He, legend shown at the bottom of the figure). The genetic differentiation between each population pair is shown with a heatmap for each species. Brighter colours in the heatmap represent higher genetic differentiation (Fst) between populations. Average fruit length for each species is also indicated.

Genomic DNA was extracted using the DNeasy Plant Kit (Qiagen) following the manufacturer’s instructions. DNA was quantified using a Qubit 4.0 fluorometer (Invitrogen), and five to seven individuals per population per species that contained the highest DNA concentration were selected for genomic library preparation. DNA samples were processed following a modified version of the double-digest restriction-site-associated DNA sequencing (ddRAD) protocol (Peterson, Weber, Kay, Fisher, & Hoekstra, 2012). For each selected individual, 200 ng of DNA was used and digested with two restriction enzymes, EcoRI and MspI. Then, barcodes were ligated and 48 individuals were pooled per library and purified with the Promega-Wizard Kit. This was followed by size selection on a 2% agarose gel using Pippin Prep electrophoresis (Sage Science) to obtain fragment lengths between 350 and 450 bp.

Subsequently, Polymerase Chain Reaction (PCR) was used to increase the number of fragments by performing 24 individual PCRs for each pool (12 cycles each), that were later combined and purified using Promega-Wizard Kit and AMPure beads. The quality of each library was controlled on a Bioanalyzer 2100 (Agilent Technologies). The libraries were sequenced with a unique barcode and a unique index per individual in paired end (PE 150bp) using an Illumina HiSeq 2000 at the Helmholtz Centre for Environmental Research – UFZ, Department Computational Biology, Leipzig, Germany.

### ddRAD data filtering and SNP calling

Raw sequence demultiplexing, quality filtering, and genotyping was performed using the software Stacks v2.53 (Catchen, Amores, Hohenlohe, Cresko, & Postlethwait, 2011, Rochette, Rivera-Colón, & Catchen, 2019) on the EVE High Performance Computing platform at the Helmholz Centre for Environmental Research, Leipzig. Demultiplexing was performed by running *process_radtags*, removing reads with uncalled bases or low-quality scores, and rescuing barcodes and RAD-tags (-clean, -quality, -rescue). From this step onwards, all analyses were carried out for each species independently. Stacks’ parameters (-M, -n, -m) were optimized per species following the r80 method by Paris, Stevens, & Catchen (2017) with the denovo_map.pl program from Stacks v2.53. More detailed information on parameter settings for each species can be found in Appendix S1 from supporting information.

We then ran the *populations* program with the population map which identifies each individual sample with a natural population (sampling site) (Figure 2, Figure S1). To control for false-positive loci, we ran populations with -r 0.75 (a locus must be found in 75% of individuals of a single population to be processed), -min-maf 0.05 (a variable site must possess a minimum minor allele frequency of 5% to be included) --max-obs-het 0.7 (a variable site must have observed heterozygosity <0.7 to be included) and --write-single-snp (restricting data analysis to the first SNP per locus). We used VCFtools/0.1.16 to explore the data for variant mean depth (--site-mean-depth) and missing data per individual (--missing-indv). To mitigate missing data and recover a higher number of loci, we followed the protocol of Cerca et al. (2021) and deleted ‘bad apples’ (i.e., individuals with more than 90% missing data) from the population map (Figure S1, Table S2), and reran the *populations* program. Then, we further filtered the data with VCFtools to remove indels (--remove-indels), retaining only sites with less than 50% missing data (--max-missing 0.5), retaining only biallelic SNPs (--min-alleles 2, --max-alleles 2), with a minimum mean read depth of 5 and maximum mean depth of 100 (--min-meanDP 5, --max-meanDP 100, --minDP 5, --maxDP 100). This approach allowed us to be more selective about the retained loci and SNPs required for model assumptions in downstream analyses, and also to recover a higher number of loci after the removal of the ‘bad apples’ (Table S3).

### Genetic diversity and genetic differentiation of palm populations

Genetic diversity and genetic differentiation parameters - including population level expected (He) and observed (Ho) heterozygosity, nucleotide diversity (π) and pairwise population differentiation (Fst) were calculated using the *populations* program in Stacks v2.53 with the -- fstats option.

### Population structure and evolutionary relationships of palm populations

We assessed population structure within species using ADMIXTURE v.1.3.0 (Alexander et al., 2009). To run ADMIXTURE, the vcf file created by the *populations* program from Stacks was modified with a custom R script (R version 4.1.2; R Core Team, 2021) to use with PLINK (by transforming chromosome from numeric to non-numeric). This modified vcf file was fed to PLINK v.1.90 (Purcell et al., 2007) to obtain a bed file, which was used to run ADMIXTURE. ADMIXTURE was ran assuming 1-10 genetic clusters (K), with a total of 5 replicates for each K, and we determined the best K by estimating the 10 fold cross-validation error.

To explore evolutionary relationships between all individuals and populations of each species, we reconstructed a phylogenetic tree for each species. Phylogenetic reconstruction was based on the maximum likelihood (ML) method using RAxML v.8.2.12 (Stamatakis, 2014). A de novo alignment was created with the *populations* program from Stacks, using a population map as input file with each individual assigned to a different population (as many populations as individuals), and with the option --phylip-var. This created an alignment including only variable sites, which was the input file for the RAxML software. We used the “-f a” and the “autoMRE” options, which determine when enough bootstrap analyses have been performed to reach convergence, using the majority-rule consensus tree criterion. We used the GTRGAMMA model of rate heterogeneity and the Lewis correction for SNP ascertainment bias (Tamuri & Goldman, 2017). The best tree for each species was visualized in R and rooted with function *midpoint*.*root* from package ‘phytools’ (Revell, 2012).

### Explanatory variables for palm population genetics

Variation in genetic diversity and genetic differentiation among palm populations can be explained by a range of variables. To test our hypotheses, we focused on frugivory-related variables (extant frugivores, extinct frugivores), human-related variables (road density, human population density) and we corrected for the potential confounding effect of the abiotic environment (temperature seasonality, precipitation seasonality, proximity to rivers and forest cover; Riordan et al., 2016; Andriamihaja et al., 2021). Although we considered additional abiotic variables (e.g., mean annual precipitation, mean annual temperature, soil characteristics), these were highly correlated to the selected variables, and therefore removed to avoid over-parameterization in the models.

Climatic variables were retrieved from https://www.worldclim.org/ (1 km resolution), rivers from http://landscapeportal.org/, percentage of forest cover for 2010 from https://madaclim.cirad.fr/ and human-related variables (e.g., road density and human population density from 2010) from Venter et al. (2016) (1 km resolution). To assess co-occurrence between palm populations and frugivores, we focused on the 93 Malagasy frugivore species identified in Méndez et al. (2022), and used the current polygon ranges from the International Union for Conservation of Nature, ver. 2020-2 (IUCN 2020). Similarly, for the extinct megafrugivores, we used the reconstructed historical distribution maps from Méndez et al. (2022), including 14 species of megafrugivores, which were modelled using co-occurrence of extant and extinct taxa across fossil sites. For extinct giant tortoises, we used the historical ranges in Pedrono et al. (2013). All climatic variables, human-related variables, and frugivore species identities, were extracted for each of the 25 palm populations with function *extract* from the ‘raster’ (Hijmans, 2020) R package. For each population, the minimum distance to a river was calculated with function *gDistance* from the R package ‘rgeos’ (Bivand & Rundel, 2021). Furthermore, we calculated the extant and extinct frugivore richness for each palm population, and identified the number of shared extant and extinct frugivore species for each population pair. Similarly, because Fst values assess population differences (as a distance matrix), we calculated population distances (differences) for each climatic variable, using function *dist* from R package ‘vegan’ (Oksanen et al., 2020). For rivers, forest cover, road density and human population density, we calculated the sum of the variable for each population pair, to account for connectivity and human impact (i.e., higher sums reflect higher potential for connectivity through more available rivers, forests, roads or humans).

### Species distribution models for palms

To predict the potential current distribution of the palm species, and therefore the potential connectivity between their populations, we constructed species distribution models for each of the four species (SDMs; Araújo et al., 2019) using the R package ‘sdm’ version 1.1-8 (Naimi & Araujo, 2016) and the workflow from Barratt et al. (2021). We used the assembled present-day palm species occurrence data from Méndez et al. (2022) and also included species records from GBIF (Global Biodiversity Information Facility) to improve accuracy of the niche models by maximising the number of unique occurrences. After cleaning the data by visually identifying errors and removing duplicates, the dataset included 103 occurrence records for *B. madagascariensis*, 191 occurrence records for *B. nobilis*, 193 occurrence records for *H. coriacea* and 310 occurrence records for *C. madagascariensis*. Since species occurrence data tends to be clustered due to sampling bias, we reduced the spatial clustering of the occurrence data by spatially thinning it (removing occurrences that are closer than 3 km to each other) with the ‘spThin’ R package (Aiello-Lammens et al., 2015). After thinning the data, the final dataset included 37 occurrence records for *B. madagascariensis*, 48 records for *B. nobilis*, 64 records for *H. coriacea* and 80 records for *C. madagascariensis* (Figure S2).

We made an *a priori* selection of 22 environmental predictors that we considered as potentially important for palm distributions (see Appendix S2 in supporting information). To avoid collinearity between predictors, we used function *vifstep* from R package ‘usdm’ (Naimi et al., 2014), which calculates the variance inflation factor (VIF) for the group of predictors, and excludes strongly correlated variables (VIF>10) in a stepwise procedure. This procedure was repeated for each species, resulting in 10 predictors for *B. madagascariensis* and *B. nobilis*, and 14 predictors for *H. coriacea* and *C. madagascariensis* (for specific predictors per species see Appendix S2 in supporting information). To generate habitat suitability maps, we constructed an ensemble model for each species based on five commonly used species distribution modelling methods: generalized linear models (Nelder & Wedderburn, 1972), generalized additive models (Hastie & Tibshirani, 1990; Wood, 2011), boosted regression trees (Friedman, 2001; Miller, Lubke, McArtor, & Bergeman, 2016), random forests (Breiman, 2001; Liaw & Wiener, 2002), and maximum-entropy (Hijmans, Phillips, Leathwick, & Elith, 2021; Phillips, Aneja, Kang, & Arya, 2006) implemented in the ‘sdm’ R package, with function *sdm*. To generate background points (pseudo-absences) that sufficiently represent the available environmental and geographic space for each species, we first created a 100 km buffer around presence points, and then generated 1000 random points inside that buffer (Figure S2; VanDerWal et al., 2009). To evaluate the accuracy of the models, they were calibrated on 30% of the data and evaluated on the remaining 70% using the ‘subsampling’ partitioning method, and by repeating each model five times. The final ensemble models were built by weighted-averaging the individual models proportionally to their AUC and TSS scores, only including models that performed adequately (AUC > 0.8, and TSS > 0.5; Swets, 1988) with function *ensemble*.

### Isolation by distance and isolation by resistance between palm populations

Isolation-by-distance (IBD – which predicts an increase of genetic differentiation between populations with increasing geographical distance) and isolation-by-resistance (IBR – which also takes potential “barriers” to gene flow into account when calculating the “shortest” [least resistant] distances between populations) may provide a more neutral expectation for genetic differentiation across Malagasy palms – regardless of dispersal by past or present frugivores or humans. To estimate the degree of connectivity between populations for each species, and thus IBR, we used the current climate suitability maps resulting from the species distribution models, and measured IBR based on least-cost distance. That is, the distance between two populations with least-accumulative costs, where cost is a function of the probability of occurrence of the species. For details on the procedure of calculating least-costs paths between all populations see Appendix S3 in supporting information.

To evaluate whether IBD or IBR explained genetic differentiation in our four palm species, we performed Mantel tests (Mantel, 1967) between all pairs of populations for each species (three population pairs for *B. madagascariensis*, 21 for *H*. coriacea, 28 for *B. nobilis* and 21 for *C. madagascariensis*), using genetic distance calculated as [Fst/(1 − Fst)] (Rousset, 1997) as the response (function *mantel*.*rtest* from R package ‘ade4’, 10,000 permutations; Bates et al., 2015).

### Determinants of palm genetic diversity and genetic differentiation

To evaluate whether dispersal limitation due to the extinction of megafrugivores has led to lower genetic diversity and higher genetic differentiation in species with megafruits compared with the small-fruited palm species, or, alternatively, that species with megafruits have not (yet) suffered from dispersal limitation, we first performed a one-way analysis of variance (ANOVA), with R function *aov*. We coupled this with a pairwise Tukey post-hoc test of honestly significant differences (R function *TukeyHSD*) in which population-level estimates of genetic diversity and genetic differentiation were compared for significant differences between the three fruit size classes: large megafruits (*B. madagascariensis*), medium megafruits (*H. coriacea* and *B. nobilis*) and small fruits (*C. madagascariensis*).

Furthermore, we used linear mixed-effects models to identify the main dispersal-related, human-related and abiotic variables associated with genetic diversity and genetic differentiation across all palm populations. We fitted full models, including all predictor variables, under the maximum-likelihood algorithm with each population-level genetic parameter as a response variable (four models in total: He, Ho, π and Fst). All climatic, frugivory-related and human impact variables were included as fixed effects, and ‘species’ as a random effect. Specifically, for genetic diversity (He, Ho and π) as the response variable, we evaluated the (fixed) effects of extant frugivore richness, extinct megafrugivore richness, road density, human population density, temperature seasonality, precipitation seasonality, distance to the nearest river and forest cover. For genetic differentiation (Fst) as the response variable, we evaluated whether the number of shared extant and extinct (mega)frugivores between populations, cumulative human population density, road density, proximity to rivers, forest cover and differences in temperature seasonality and precipitation seasonality explained variation in Fst across palm populations. To assess differences between palms with megafruits and the small-fruited palm, we repeated the Fst model by only including the palm species with megafruits. Such a subset analysis was not possible for the small-fruited palm due to small sample size. Similarly, an analysis only including species with megafruits for the genetic diversity analysis was not possible due to small sample size.

Mixed-effect models were fitted with function *lmer* from R package ‘lme4’. All possible models were evaluated with function *dredge* from R package ‘MuMIn’ (Barton, 2022). This included the evaluation of 256 models for each response variable. To account for uncertainty and model selection bias, instead of selecting only the ‘best model’, we carried out multimodel selection and model averaging based on the Akaike information criterion (AICc) and retained a set of top models (‘best models’) with ΔAICc ≤ 2 from the ‘best’ model (Grueber et al., 2011). Coefficient estimates for each predictor on the response were calculated using the ‘conditional’ average, which only averages over the models that included the predictor.

## Results

### ddRAD data filtering and SNP calling

We obtained paired-end Illumina reads for 28 individuals of *B. madagascariensis* (228464 retained reads), 47 of *H. coriacea* (513399 retained reads), 63 of *B. nobilis* (866044 retained reads) and 46 of *C. madagascariensis* (665801 retained reads, Table S3). Stacks output haplotype files contained between 5,091 (*B. madagascariensis*) and 26,435 (*C. madagascariensis*) variant sites (Table S3). After filtering for ‘bad apples’ (i.e., removing individuals with more than 90% missing data) one individual of *B. madagascariensis*, two of *H. coriacea*, 11 *B. nobilis*, and one of *C. madagascariensis* were removed (Table S2). After excluding non-bi-allelic loci, filtering for minor allele frequencies, maximum observed heterozygosity, and excluding sites with more than 50% missing data, final numbers of SNPs per species used for subsequent analyses (Table S3) were as follows: 1,926 (*B. madagascariensis*), 12,898 (*H. coriacea*), 5,519 (*B. nobilis)* and 11,589 (*C. madagascariensis*). The demultiplexed ddRAD sequences are available from ENA (European Nucleotide Archive) project PRJEB56299.

### Genetic diversity and genetic differentiation between fruit size classes

We did not find support for the hypothesis (H1) that dispersal limitation has led to low genetic diversity and high genetic differentiation in megafruit species (Figure 1). Instead, per population genetic diversity (Figures 2 and 3) was significantly higher for palms with megafruits (*B. madagascariensis, H. coriacea* and *B. nobilis*) for all three metrics, than for the small-fruited palm (Figure 3, ANOVA F-value = 3.77, p < 0.05, Table 1, Table S6). Average Ho was highest for *B. madagascariensis* (Ho=0.245), closely followed by *H. coriacea* (Ho=0.244) and *B. nobilis* (Ho=0.223) and significantly lower for *C. madagascariensis* (Ho=0.193). However, π and He were highest for *H. coriacea* (π=0.259; He=0.232), followed by *B. madagascariensis* (π=0.250; He=0.230) and *B. nobilis* (π=0.231; He=0.206), but again lowest for the small-fruited *C. madagascariensis* (π=0.198; He=0.179). For detailed information on population-level values of all the genetic diversity metrics see Table S7.

**Figure 3.**
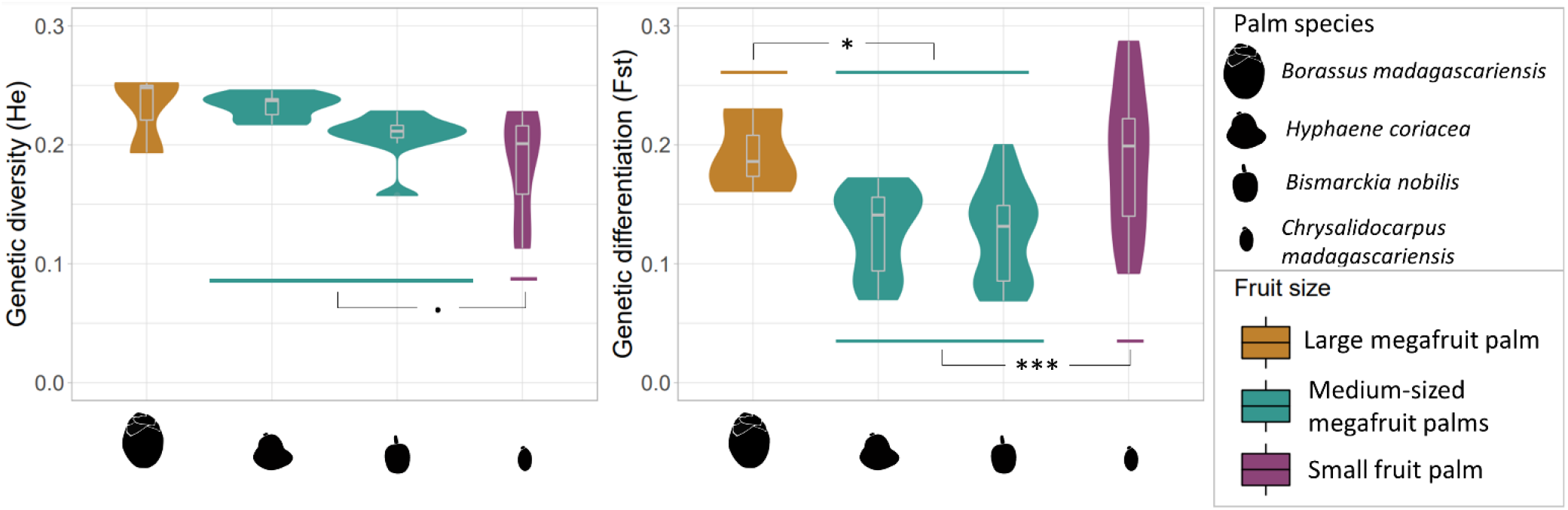
Violin plots with inner boxplots showing the genetic diversity (left - He) and genetic differentiation (right - Fst) per species within each fruit size class: large megafruits (dark yellow), medium-sized megafruits (green) and small fruits (purple). Results from the Tukey honest significant test are shown: ***p<0.001, *p<0.05, **·**p=0.05.

Similarly, for genetic differentiation, the highest genetic differentiation (Fst) was found among populations of the small-fruited palm (Figures 2 and 3), but the variance in Fst was also high for this species (Table S8). The Fst estimates from the small-fruited palm did not differ significantly from those of the largest-fruited *B. madagascariensis*, which also showed high levels of genetic differentiation (Figures 2 and 3, ANOVA F-value = 15.37, p < 0.001, Table S6 and Table S8). Significantly lower Fst values were found between populations of palms with human-used, medium-sized megafruits (*B. nobilis, H. coriacea)* compared to the other two palm species (Figures 2 and 3, Table S6).

### Population structure and evolutionary relationships of palm populations

The results from ADMIXTURE analyses showed evidence for a generally low intraspecific genetic structure across species (Figure 4), with k=2 genetic clusters for all four species. In all species, the two clusters were generally consistent with the geographical separation of northern and southern populations. In *B. madagascariensis* and *C. madagascariensis*, results of cross validation error between k=2, k=3 and k=4; and k=2 and k=3, respectively, were very close and it was therefore difficult to conclude the most appropriate k value for these two species (see Figures S3 and S4).

**Figure 4.**
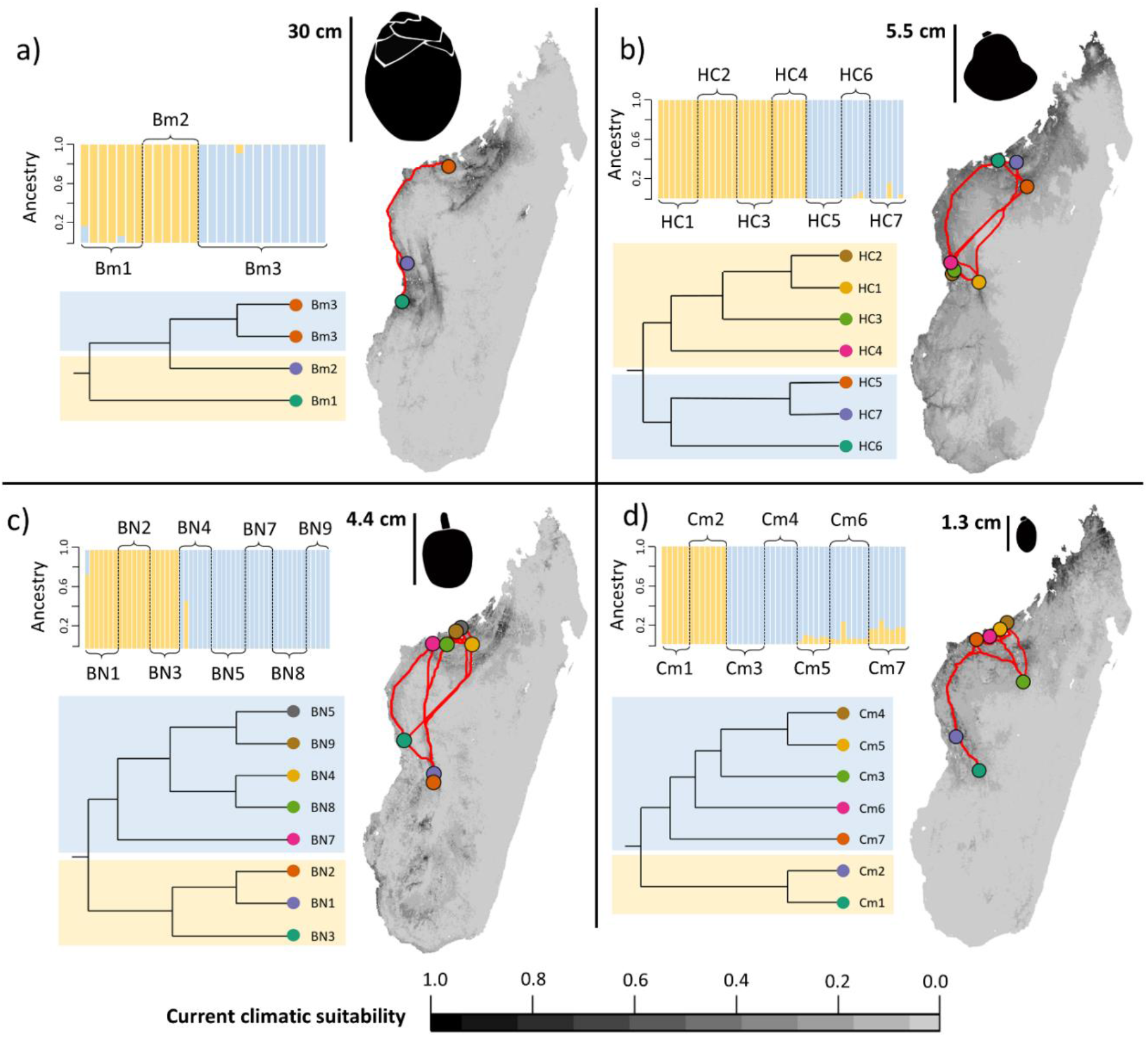
Sampled populations against the current climate suitability maps from ensemble species distribution models for four Malagasy palm species. Darker colours indicate more suitable areas (legend shown at the bottom of the figure). The reconstructed phylogenetic trees among populations are shown for each species. Each population has a unique colour that matches both the phylogeny (tip) and sampling locality on the map. The background colours in the phylogenetic trees are referring to the two genetic clusters which are also shown with the admixture bar plots per species. Each horizontal bar represents a different individual, with the identified admixture proportions among the clusters. Clusters reflect the north/south separation of populations: the “southern” group of populations in yellow and the “northern” group of populations in blue. Finally, the red lines between populations on the map represent least-cost distances between population pairs based on the climatic suitability. a) *Borassus madagascariensis*, b) *Hyphaene coriacea*, c) *Bismarckia nobilis* and d) *Chrysalidocarpus madagascariensis*. Average fruit length for each species is also indicated.

The phylogenetic reconstruction showed additional information on genetic structuring of individuals into clades, or populations, generally supporting clades of individuals sampled from the same ‘natural’ population (Figure 4, see Figure S5 for the full phylogenetic trees). Interestingly, for *B. madagascariensis*, the phylogeny showed a separation of population Bm3 into two additional clades, even though these individuals were only 2 km apart (Table S1, Figures 4 and S5a), in line with the results from ADMIXTURE with cross validation errors very close between k=2, k=3 and k=4 (separating population Bm3 into two clusters, Figure S4). This indicates higher genetic structure for *B. madagascariensis*, since we sampled 3 ‘natural’ populations and the number of clusters goes from k=2-4, showing high genetic structuring among the 3 populations. In *H. coriacea* both admixture and phylogenetic tree supported two populations/clades, with population HC6 being ancestral for the northern clade, and population HC4 ancestral for the southern clade (Figures 4 and S5b). Similarly, in *B. nobilis*, the north-south differentiation was recovered with population BN3 ancestral in the south, BN7 in the north, but not all individuals clustered within a clade that reflected their natural population, suggesting high admixture between populations (especially for BN1-BN2 and BN4-BN8, Figure S5c). Finally, for the small-fruited *C. madagascariensis*, besides clear genetic differentiation between northern and southern populations, population Cm3 was also recovered as a separate cluster when considering k=3 in the admixture analysis. This population is located geographically between the northern and southern populations and therefore could be the reason to be found as a separate cluster (Figure 4, Figures S4 and S5d).

### Habitat connectivity and geographical distance effects on genetic differentiation

Species distribution models generally showed good performance (AUC > 0.8, and TSS > 0.5; Figure 4, Table S4). Mantel tests showed that both IBD and IBR significantly explained genetic differentiation among populations for all species (p-value <0.05), except for *B. madagascariensis*, probably because of the small sample size for this species (Table S5, Figure S6). For *H. coriacea* (IBD Mantel-r = 0.886, IBRr = 0.877), *B. nobilis* (IBDr = 0.803, IBRr = 0.841), and *C. madagascariensis* (IBDr = 0.777, IBRr = 0.864), IBD and IBR were strongly correlated with normalized genetic differentiation (Fst, Figure 2), illustrating that distance between populations also often reflects environmental resistance (Figure 4). These results illustrate that both geographical distance and landscape resistance explain variation in genetic differentiation across palm populations in Madagascar.

### Determinants of palm genetic diversity and genetic differentiation

Selection of the best mixed-effect models (models with ΔAICc ≤ 2 from the best model) resulted in three models for He and Ho, four models for π, and two models for Fst when including all palms, and four models for Fst when only including the three megafruit palm species (Table S9).

Interestingly, our results suggest that historical long-distance dispersal events by megafrugivores are still reflected in the genetic differentiation of palm populations, thus consistent with H2. Specifically, the number of shared past (now extinct) megafrugivores was significantly negatively correlated with genetic differentiation (Fst) between populations (Figures 5 and 6, Table S9), suggesting that historical long-distance dispersal by megafrugivore dispersers may have increased gene flow between populations that shared megafaunal interaction partners. In comparison, shared extant frugivores were not significantly correlated with Fst among populations (Figure 6). Results for genetic diversity did not support H2.

**Figure 5.**
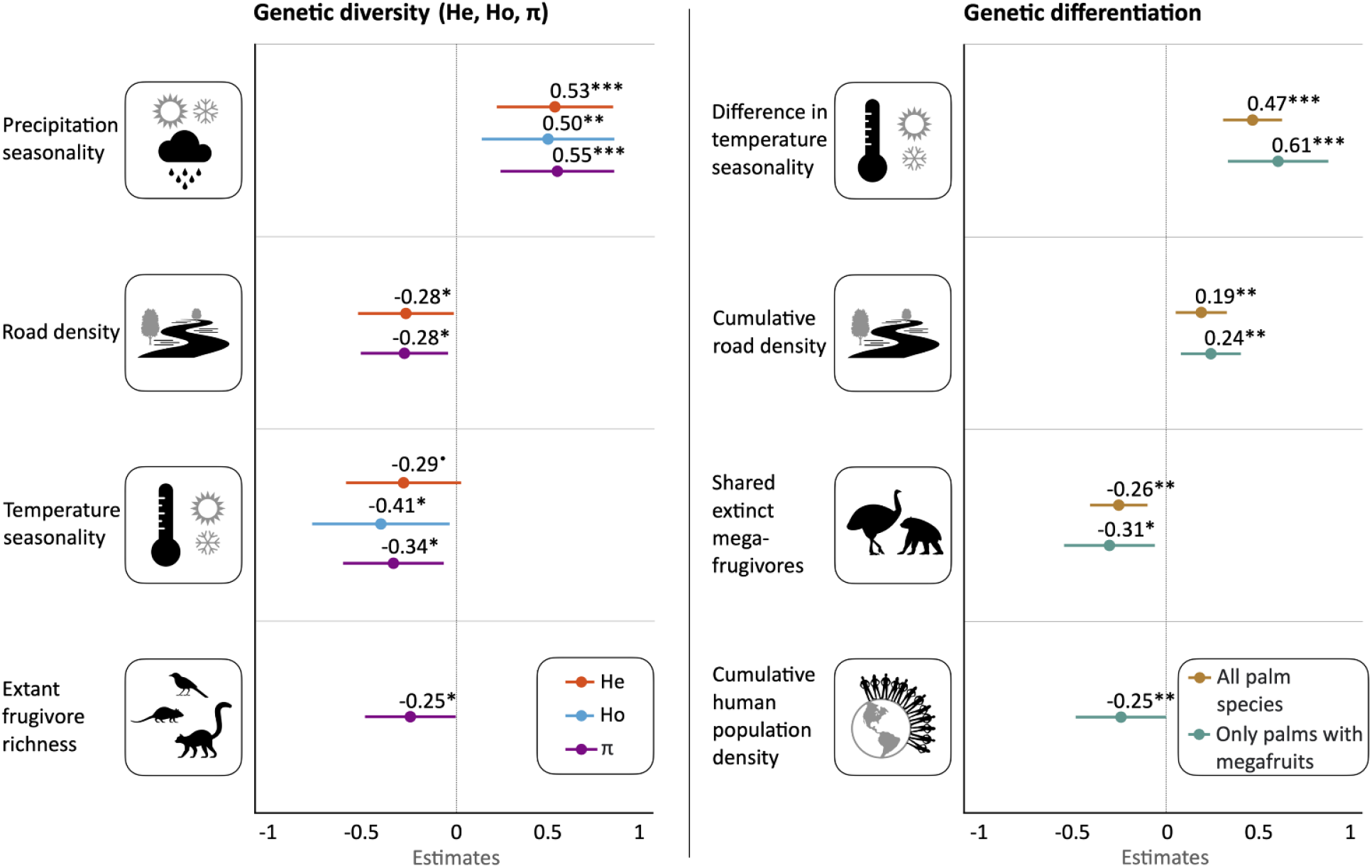
Results from the linear mixed effect models showing the standardized effects of predictor variables on the genetic diversity (left) and genetic differentiation (right) of palm populations in Madagascar. Only significant predictors after model averaging are shown. The average conditional estimates (circles) are shown by the numbers next to each bar, with the 95% confidence intervals (bars) of those estimates. Significance of each predictor variable is also given ***p ≃0; **p < 0.001; *p < 0.01; **·**p=0.05. See Table S9 for detailed results of the models.

**Figure 6.**
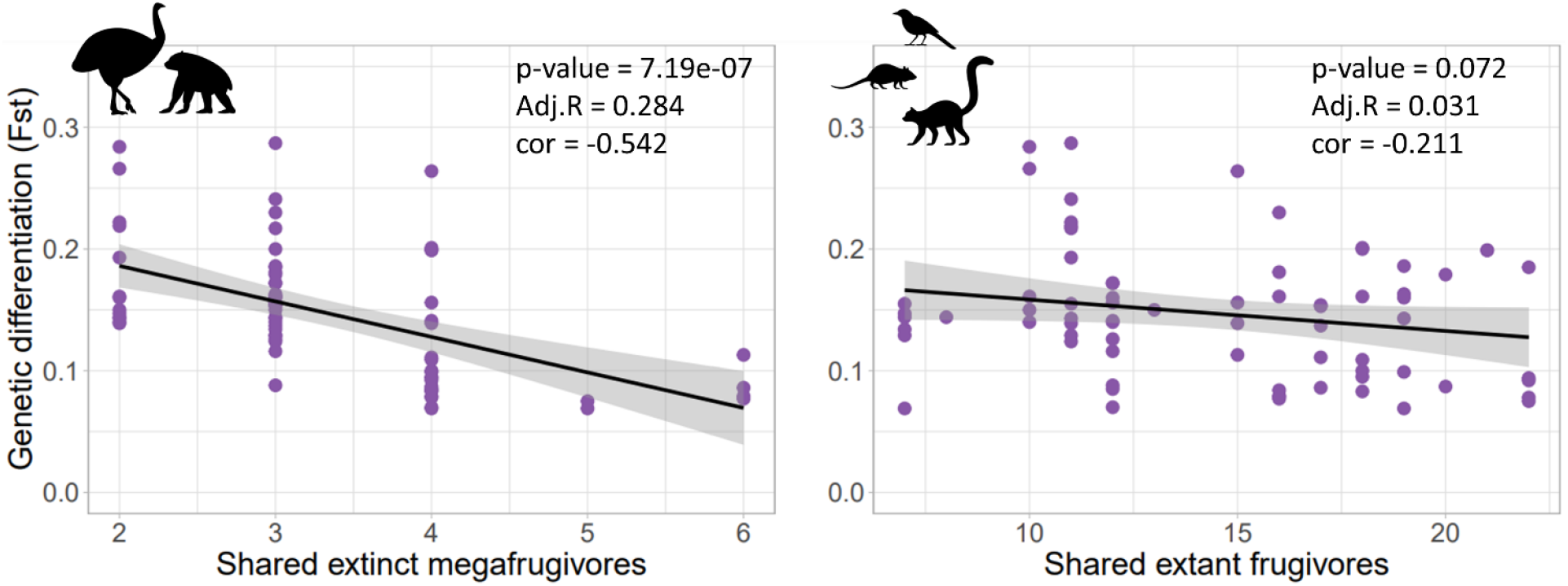
Relationship between the genetic differentiation (Fst) between all (within species) population pairs across four Malagasy palm species and the number of shared extinct megafrugivores (left) and number of shared extant frugivores (right). Only the relationship with the extinct megafrugivores was statistically supported in the linear mixed-effects models (see Figure 5). Showing statistical values from a simple linear model; p-value, Adj.R= adjusted R square; cor = correlation value.

Specifically, extinct megafrugivore richness did not explain variation in genetic diversity across palm populations, while extant frugivore richness had a significant negative relationship with π. We found mixed support for H3. Road density had a positive correlation with Fst (Figure 5), suggesting that higher human impact reduces, rather than increases, gene flow when including all four palm species. This is consistent with results for genetic diversity, which indicated lower genetic diversity with higher road densities (for Ho and π, not for He, Figure 5). Nevertheless, the Fst model including only population pairs of palms with megafruits indicated a significant negative correlation between human population density and Fst (Figure 5, Table S9). This suggests that higher human population density may also lead to genetically more similar palm populations, thus supporting a potential human dispersal role for palms with megafruits (H3).

Finally, differences in temperature seasonality between populations was the strongest determinant of Fst, and higher temperature seasonality was associated with lower genetic diversity (Figure 5, Table S9). In addition, higher precipitation seasonality was associated with higher genetic diversity (Figure 3, Table S9). Proximity to rivers and forest cover were not significantly associated with genetic diversity or genetic differentiation.

## Discussion

Contrary to our expectation, we did not find evidence for a loss of genetic diversity or higher genetic differentiation in palms with megafruits due to the loss of their megafrugivore dispersers (Figures 1 and 3). In fact, genetic diversity was higher in species with megafruits than in the species with small fruits, and genetic differentiation did not differ significantly between the largest- and smallest-fruited species (Table 1, Figure 3). Instead, the legacy of megafrugivores regularly achieving long dispersal distances is still reflected in the genetic diversity and genetic differentiation of palms that were formerly dispersed by such large animals. Specifically, a higher number of shared extinct megafrugivore species between palm population pairs was associated with lower Fst values, indicating higher gene flow, whereas no relationship with extant frugivores – that are not able to swallow and disperse the seeds – was found (Figure 6). Furthermore, genetic diversity increased with fewer roads, illustrating the potential negative effect of human-driven fragmentation on palm genetics (Figure 5). However, genetic differentiation was lowest for the human-used medium-sized megafruit species, and decreasing genetic differentiation between populations was associated with increasing human population density, but only for species with megafruits (Figure 3 and 5). This suggests a potential contribution of human-mediated dispersal to gene flow in large-fruited, human-used Malagasy palms. Finally, besides frugivory- or human-related dispersal, palm genetic diversity and genetic differentiation were strongly linked to abiotic and geographical gradients (Figures 2, 4 and 5). For example, broad-scale genetic structure of western Malagasy palms is shaped by isolation-by-distance and isolation-by-resistance, leading to a north / south separation of populations (Figures 4 and S5).

### Methodological issues

Genetic diversity and genetic differentiation are complex measures subjected to a myriad of ecological and evolutionary processes. Besides gene flow, genetic diversity and genetic differentiation in plants are the result of demographic history of the species (population size over time), selection, drift and mutation (Ellegren & Galtier, 2016). Although our results are consistent with some of our expectations (Figure 1), our sample size (four species, 25 populations) is limited. Furthermore, it is difficult to exclude the possibility that other co-varying or confounding variables, such as pollination, life history, and sexual system, may have influenced patterns of genetic diversity and genetic differentiation. Nevertheless, the use of genome-wide genetic data, a comparative framework to evaluate our hypotheses, and robust statistical analyses, provides at least a solid foundation to evaluate questions at the interface of population genomics and ecology.

### No signature of megafrugivore extinctions in Malagasy palms with megafruits

In contrast to our expectation (H1), species-level genetic diversity increased with fruit size, whereas genetic differentiation did not significantly differ between the largest- and smallest-fruited palm (Figures 1 and 3). In line with this result, a meta-analysis including 102 Neotropical plants indicated no difference in genetic diversity of species dispersed by megafrugivores compared to non-megafrugivore dispersed species, even though almost all megafauna was lost in the Neotropics during the Late Quaternary extinction wave (Collevatti et al., 2019). Extant frugivores and secondary seed dispersers may counterbalance the seed dispersal loss due to Quaternary megafrugivore extinctions, thus possibly maintaining genetic diversity in these megafruit plant species. In addition, pollination may contribute to gene flow between plant populations (Collevatti et al., 2019). This may also apply to Malagasy plant species with megafruits, although evidence for secondary seed dispersal (e.g., by rodents) in Madagascar is scarce (but see Méndez et al., 2022; Razafindratsima, 2014).

The relatively high genetic diversity of our western Malagasy palm species with megafruits is consistent with findings from other Malagasy palms. For example, in the eastern humid forests of Madagascar, the palms *Lemurophoenix halleuxii* and *Voanioala gerardii* also bear megafruits, and have relatively high within-population genetic diversity (Shapcott et al., 2012). However, in contrast to the western savanna palms (at least *H. coriacea* and *B. nobilis*) these rainforest species have small, fragmented populations, and are (critically) endangered (IUCN, 2012). Furthermore, the more widespread small-fruited *Dypsis decipiens* has lower genetic diversity than the narrow-ranged, small population (ca. 100 individuals left) of small-fruited *Dypsis ambositrae* (Gardiner et al., 2017), illustrating how fruit size, and potential frugivore-mediated dispersal, is certainly not the only determinant of plant genetic diversity. Instead, it is possible that increased human activity, habitat fragmentation and demographic change have led to the loss of genetic diversity and narrow ranges in several Malagasy palms (Helmstetter et al., 2021).

### Historical long-distance dispersal events by megafrugivores have increased gene flow in Malagasy palms

Our results support the hypothesis (H2) that long-distance dispersal events provided by megafrugivores in the past may have increased gene flow, and therefore decreased genetic differentiation and genetic structure of populations of palms (Figures 1-6). This is evidenced by the negative correlation between the number of shared megafrugivore species (i.e., giant lemurs, elephant birds, giant tortoises) between population pairs and genetic differentiation (Figures 5 and 6). Increased gene flow in the past may also explain the higher genetic diversity in palms with megafruits (Figures 2 and 3). Although the correlation between shared megafauna and genetic differentiation is striking, it is possible that past megafaunal distributions – which were inferred from fossil co-occurrences with extant species and their current distributions (Méndez et al., 2022) – co-vary with distance between population pairs, climate, and human settlement, i.e., factors that also directly interact with palm genetics (Figures 4 and 5). Nevertheless, the comparison with extant frugivores (Figure 6), which would be subjected to similar environmental correlations but did not show a significant relationship with genetic differentiation, provides some evidence for the interpretation that past dispersal events have increased gene flow between palm populations, and the loss of these dispersal services by megafrugivores has not yet manifested itself in population genetic differentiation. Our results contrast with defaunation and downsizing of frugivorous lizards in the Canary Islands, which led to reduced population connectivity and high genetic differentiation among populations (Pérez-Méndez et al., 2018), thus illustrating the expected effect of defaunation on genetics. There are two possible explanations why our results did not show this same pattern.

First, it is possible that gene flow between Malagasy palm populations has been maintained by other means of seed dispersal such as barochory, hydrochory and secondary seed dispersal (Blanco et al., 2019), or pollen dispersal (Giombini et al., 2017). Furthermore, human use, and thereby (occasional) dispersal of fruits, may have facilitated gene flow between palm populations (H3, Figure 1). Indeed, higher human population density was associated with lower genetic differentiation among populations of palms with megafruits (Figure 5), and the palms showing the lowest genetic differentiation and highest admixture (low genetic structure) (Figures 2–4) are those known to be essential to the daily life of local people in the western part of Madagascar, i.e., *H. coriacea* and *B. nobilis* (Rakotoarinivo et al., 2020). Increased human-use of the palms with megafruits (i.e., *H. coriacea* and *B. nobilis*) could have increased movement of palm material (including seeds) between sites, with possible effects on gene flow and reduced genetic differentiation in natural populations. The level of utilisation of the palms included in our study negatively correlates with their average Fst values (higher level of utilisation, lower genetic differentiation; see Rakotoarinivo et al., 2020 for level of utilisation, Figure S7). In line with our results, human-mediated dispersal (based on diet) has been shown to be the most important dispersal mode for a large proportion of large-fruited Neotropical species (that used to rely on extinct megafrugivore dispersal), leading to larger geographical range sizes of megafruit species used by humans compared to those not used by humans (Van Zonneveld et al., 2018).

Second, it is possible that our palm species do not show high levels of genetic differentiation yet, because the megafauna extinctions have been relatively recent in time (ca. 1000 years; Crowley, 2010), and genetic change may only become visible after sufficient generations. Indeed, long-living species showed the highest levels of genetic diversity across the 384 plant species assessed by de Kort et al. (2021). This ‘extinction debt’ (Tilman et al., 1994), where long-living plants respond more slowly to reduced gene flow due to their longer lifespans, may explain why these palms still show high genetic diversity and low genetic differentiation. Indeed, if we assume a generation time of c. 8-40 years for these palms (Gaut, Muse, Clark, & Clegg, 1992), this would correspond to c. 40 generations since the extinction of the megafauna, which may not be sufficient to reduce genetic diversity and increase genetic differentiation. However, Ellegren & Galtier, (2016) argued that species with longer lifespans are normally ‘K-strategists’, and therefore have lower fecundity and larger propagule sizes, which reduces their effective population size and therefore reduces their potential genetic diversity. Our results indicate higher genetic diversity and lower genetic differentiation in palms with megafruits, which may be due to a combination of factors related to alternative dispersal, life history, long lifespans and long generation times, and the detrimental effect of hindered gene flow may only become evident over time.

### Abiotic and human-related drivers of genetic diversity and genetic structure

Palm genetic diversity and genetic differentiation were also shaped by abiotic variables, such as geographic distance (IBD), environmental resistance (IBR), and climate seasonality. We found that populations with higher genetic diversity occur in areas with more precipitation seasonality, and populations with lower genetic diversity in areas with higher temperature seasonality. Meanwhile, genetic differentiation among populations was mostly driven by larger differences in temperature seasonality (Figure 5). These results may be partly explained by the ‘asynchrony of seasons hypothesis’ (Martin et al., 2009), where seasonality affects the flowering and/or fruiting timings of plant populations, leading to reproductive asynchrony, disrupting gene flow between populations and enhancing genetic differentiation. This may be even further enhanced by dioecy in the megafruit species. Furthermore, temperature and precipitation seasonality co-vary with fire activity (Saha et al., 2019), which could also facilitate isolation of populations, and thus genetic differentiation (Abrahamson, 1999).

Finally, although humans may facilitate dispersal of palm fruits, our results also show negative effects of human impact on palm genetics. Specifically, we found that road density was associated with lower genetic diversity and higher genetic differentiation (Figure 5). Roads may therefore lead to fragmentation of natural habitats and hinder gene flow. Furthermore, the western savannas of Madagascar are known to experience regular fires, mostly by regulated burnings by humans. Mature individuals of several species of palms can survive fires (Lee Arneaud et al., 2017) and *B. nobilis* and *H. coriacea* are both known to be fire resistant.

Therefore, in the case of *B. nobilis* and *H. coriacea*, human impact seems to be contributing to the expansion of their populations, specifically through fire that stimulates seed germination (Chambon et al., 2021). This could possibly lead to genetically homogeneous populations and the loss of local adaptations or rare unique alleles. Thus, human impact may have a dual effect on western Malagasy palms – contributing to the expansion as well as the fragmentation of populations. Unravelling population size changes of these palm species since human arrival and megafrugivore extinction on Madagascar may help to understand and predict how populations will behave in the face of climate change and ongoing defaunation.

## Author contributions

Laura Méndez and Renske E. Onstein conceived the ideas for the study with support from William J. Baker, Wolf L. Eiserhardt, W. Daniel Kissling and Christopher D. Barratt. Laura Méndez and Vonona Randrianasolo organized the fieldwork and obtained samples. Laura Méndez and Walter Durka conducted the laboratory work. Laura Méndez analysed the data with support from Walter Durka, Christopher D. Barratt and Renske E. Onstein. Renske E. Onstein obtained financial support for the project. Laura Méndez and Renske E. Onstein wrote the paper with reviews and editing from all other authors.

## Data Availability Statement

Demultiplexed ddRAD sequences are available in ENA (European Nucleotide Archive) project PRJEB56299. Supporting information for this study and all data used, including (mega)frugivore species lists, climatic and human-related variables per population, are openly available on the Dryad Digital Repository XXX. The code to the pipeline to conduct all the analyses of the ddRAD data is published at Zenodo (https://doi.org/10.5281/zenodo.7362068).

## Acknowledgements

We are grateful to Kew Madagascar Conservation Centre (especially Hélène Ralimanana and Stuart Cable) and the Botanical and Zoological Garden of Tsimbazaza for their invaluable help with planning and carrying out fieldwork in Madagascar. We thank Landy Rajaovelona for helping with the issuing of collecting permits 158/19/MEDD/SG/DGEF/DGRNE and 172/19/MEDD /SG/DGEF/DGRNE. We would also like to thank Hanta Razafindraibe, Henintsoa Razanajatovo, Tahina Razafindrahaja, Andry Rakotoarisoa, Roger Rajaonarison, Fidelis Randrianasolo, Angelos J. Tianarifidy, Hajatianalalaina Rakotoarimanana, Gervais (Analalava forest), Therisis and Rasolofo Kalobe (Manjato forest) for their help in the field. We thank John Dransfield, Alison Shapcott, Wolfgang Stuppy and Alexander Zizka for contributing ideas during a workshop on ‘palms of Madagascar’, and the members of the Evolution and Adaptation research group (iDiv) for additional discussions and advice. We thank Isis Petrocelli, Ina Geier, Martina Herrmann, Stephan Schreiber and Matthias Bernt for their great help during laboratory work and sequencing. We acknowledge the Helmholtz Centre for Environmental Research for providing access to EVE, the High-Performance Computer. Laura Méndez, Renske E. Onstein and Christopher D. Barratt gratefully acknowledge the support of iDiv, funded by the German Research Foundation (DFG–FZT 118, 202548816).

