## Supporting Information for "Genomic signatures of past megafrugivore-mediated dispersal in Malagasy palms"

### Table of Contents:


| | |
|---|---|
| **Appendix S1:** Parameter optimization in Stacks | Page 2 |
| **Appendix S2:** Predictor variables selection for species distribution models | Page 2 |
| **Appendix S3:** Predictor variables selection for species distribution models | Page 3 |
| **Figure S1:** Bioinformatics workflow | Page 4 |
| **Figure S2:** Occurrences and background points for each species used for the SDMs | Page 5 |
| **Figure S3:** Cross validation error per species for ADMIXTURE | Page 6 |
| **Figure S4:** Results from ADMIXTURE analyses showing K3 | Page 7 |
| **Figure S5:** Phylogenetic trees | Page 8-11 |
| **Figure S6:** Results from the mantel tests of IBD and IBR | Page 12 |
| **Figure S7:** Correlation between genetic differentiation of species and human use | Page 13 |
| **Table S1:** Coordinates of the centroids of all sampled palm populations | Page 14 |
| **Table S2**: Missing data per individual | Page 15-19 |
| **Table S3:** Number of recovered loci before and after filtering | Page 20 |
| **Table S4:** Results from SDMs | Page 21 |
| **Table S5:** Results from Mantel tests | Page 22 |
| **Table S6:** ANOVA results using Tuckey Honest Significant Difference | Page 23 |
| **Table S7:** Per population genetic diversity | Page 24 |
| **Table S8:** Genetic differentiation for each pair of populations | Page 25-26 |
| **Table S9:** Results from LMMs | Page 27 |
| **References** | Page 28 |

**Appendix S1. Parameter optimization in Stacks**

After demultiplexing the ddRAD data with *process*_radtags in Stacks v2.53 (Catchen, Hohenlohe, Bassham, Amores, & Cresko, 2013; Rochette, Rivera-Colón, & Catchen, 2019), we proceeded to optimize the parameters that need to be chosen (which significantly affect the building and quality of the resulting loci) for the *de* novo assembly following the r80 method by (Paris, Stevens, & Catchen, 2017). This step was done independently for each species. This method aims to find a balance between recovering the highest number of loci and introducing the least amount of sequencing error (finding true polymorphism). The parameters to optimize are *M* and *n. M* - number of mismatches allowed between stacks within individuals – if *M* is set too low, then some loci will fail to be reconstructed. This means the SNPs contained in that locus will not be identified and this locus will appear as two loci to the remainder of the pipeline. Setting *M* too high will allow repetitive sequence to chain together into large, nonsensical loci. *n* – number of mismatches allowed between catalog loci between individuals – if *n* is set too low, there will be loci across individuals that are represented independently in the catalog that are truly the same locus. If you set too high, again loci close together in sequence space will be allowed to chain together and create big, erroneous loci in the catalog.

Following Paris et al. (2017), we therefore ran the *denovol_map.pl* from Stacks with combinations of *M* and *n* from 1 to 9 (*M*=n) and setting *m* (depth coverage) to three, for a subset of individuals of each species separately. Then we visualized the number of polymorphic loci across 80% of the samples (r80 loci) per each *M* and *n* and how many new polymorphic loci were identified for each increment of *M* and *n*. After this we selected *M=n=4* for *B. madagascariensis*, *M*=*n*=5 for *H. coriacea*, *M*=*n*=4 for *B. nobilis* and *M*=*n*=5 for *C. madagascariensis*.

**Appendix S2. Predictor variables selection for species distribution models**

A first selection of environmental and edaphic predictors was made to avoid overparameterization. We chose 22 predictors: annual mean temperature, temperature seasonality, maximum temperature of the warmest month, minimum temperature of the coldest month, temperature annual range, annual precipitation, precipitation of the wettest month, precipitation of the driest month, precipitation seasonality (retrieved from https://www.worldclim.org/), evapotranspiration, climatic water deficit, altitude, slope, solar radiation, percentage of forest in 2010 (retrieved from https://madaclim.cirad.fr/), soil clay content, soil cation exchange capacity, soil organic carbon, pH in soil water, soil sand content, soil extractable aluminum and soil total nitrogen (retrieved from https://data.isric.org/). After extracting each predictor for each population, to avoid collinearity, we used function *vifstep* from package 'usdm' (Naimi et al., 2014), which calculates the variance inflation factor (VIF) for the group of predictors, and excludes strongly correlated variables in a stepwise procedure. After that we deleted collinear variables per species and the final predictors used per species were: 10 predictors for *B. madagascariensis* and *B. nobilis* (maximum temperature of the warmest month, annual precipitation, precipitation seasonality, slope, percentage of forest in 2010, soil clay content, soil organic carbon, pH in soil water, soil extractable aluminum and soil total nitrogen), 14 predictors for *H. coriacea* (maximum temperature of the warmest month, annual precipitation, precipitation seasonality, climatic water deficit, altitude, slope, solar radiation, percentage of forest in 2010, soil cation exchange capacity, soil organic carbon, pH in soil water, soil sand content, soil extractable aluminum and soil total nitrogen) and 14 predictors for *C. madagascariensis* (temperature seasonality, maximum temperature of the warmest month, annual precipitation, precipitation seasonality, altitude, slope, solar radiation, percentage of forest in 2010, soil cation exchange capacity, soil organic carbon, pH in soil water, soil sand content, soil extractable aluminum and soil total nitrogen).

**Appendix S3. Procedure for building least-cost-paths from ensemble species distribution models**

We used the current climate suitability maps resulting from the ensemble species distribution models, to measure IBR (Isolation-by-resistance) based on least-cost distance. A least-cost distance is the distance between two populations with least-accumulative costs, where cost is a function of the probability of occurrence of the species. To build the least-cost path, a map depicting conductance instead of resistance is needed, meaning higher values reflect higher costs for dispersal, whereas lower values reflect lower costs for dispersal. Since the niche model probability maps depicted the opposite to the conductance map (lower values – low occurrence probability – lower connectivity; higher values – higher probability of occurrence – higher connectivity), we first classified the values of each raster layer resulting from the niche models (which ranged from 0-1) into 20 categories (values from 1 to 20; function *cut* from package 'raster' with 20 breaks). Then, to have lower values (1) as lower cost and higher values (20) as higher cost, we inverted the values of each categorized raster (function *raster.invert* from package 'spatialEco', Evans, 2021). Next, we calculated a transition object for which we specified the direction of the path (the neighbourhood around each pixel for the algorithm to consider when deciding on the least-cost path from A to B). We specified the direction "queen's case", based on Moore's neighbourhood, which allows to move towards all pixels around the target pixel, even the diagonal ones (function *transition* from package 'gdistance';van Etten, 2017), and corrected the resulting transition object for map distortion with function *geoCorrection*. The resulting object was used to calculate the pairwise least-cost paths between populations for each species (function *costDistance*).

**Supporting Figures**

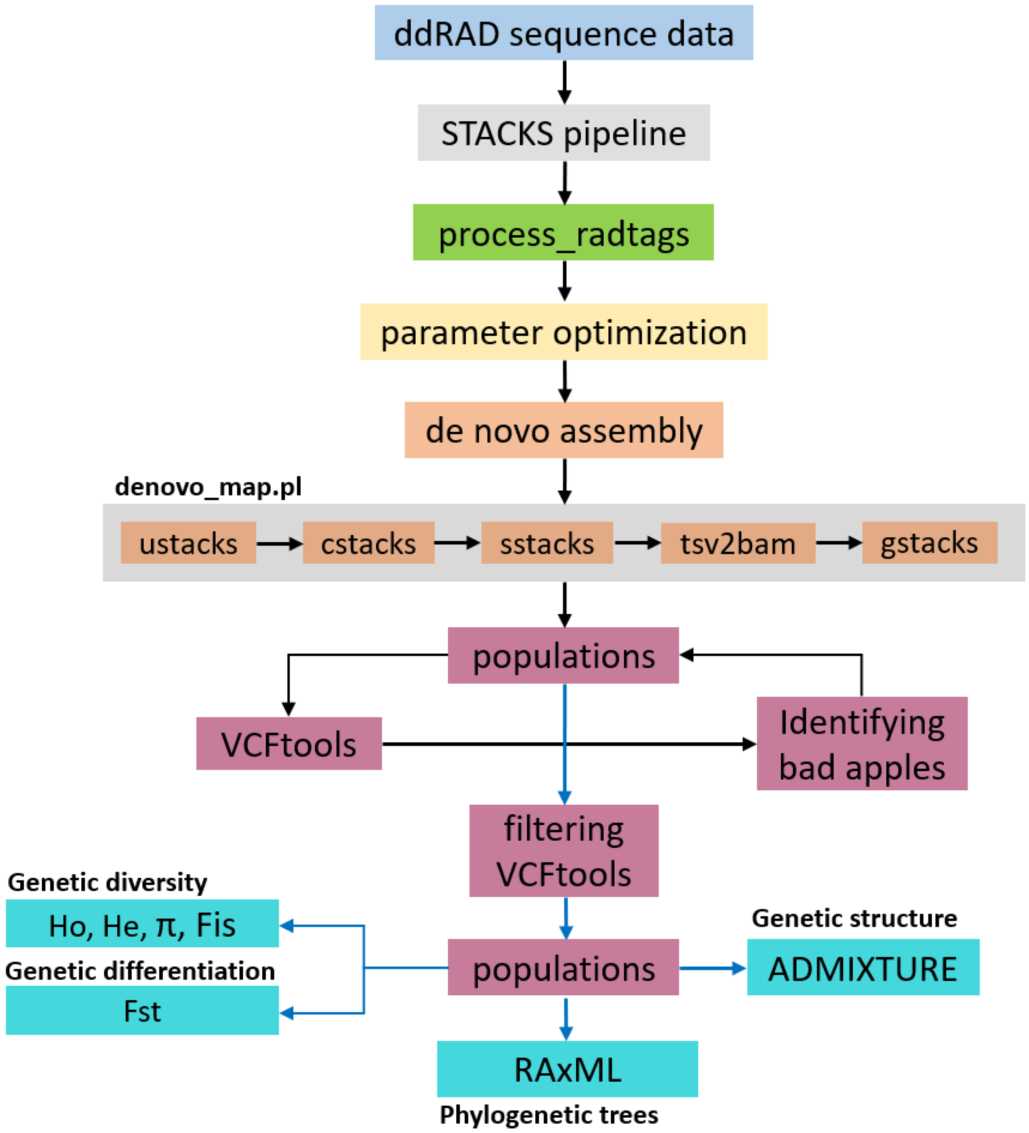


**Figure S1.** Bioinformatics workflow showing each step from the raw ddRAD sequence data to the final genetic parameters, phylogenetic trees and genetic structure analyses.

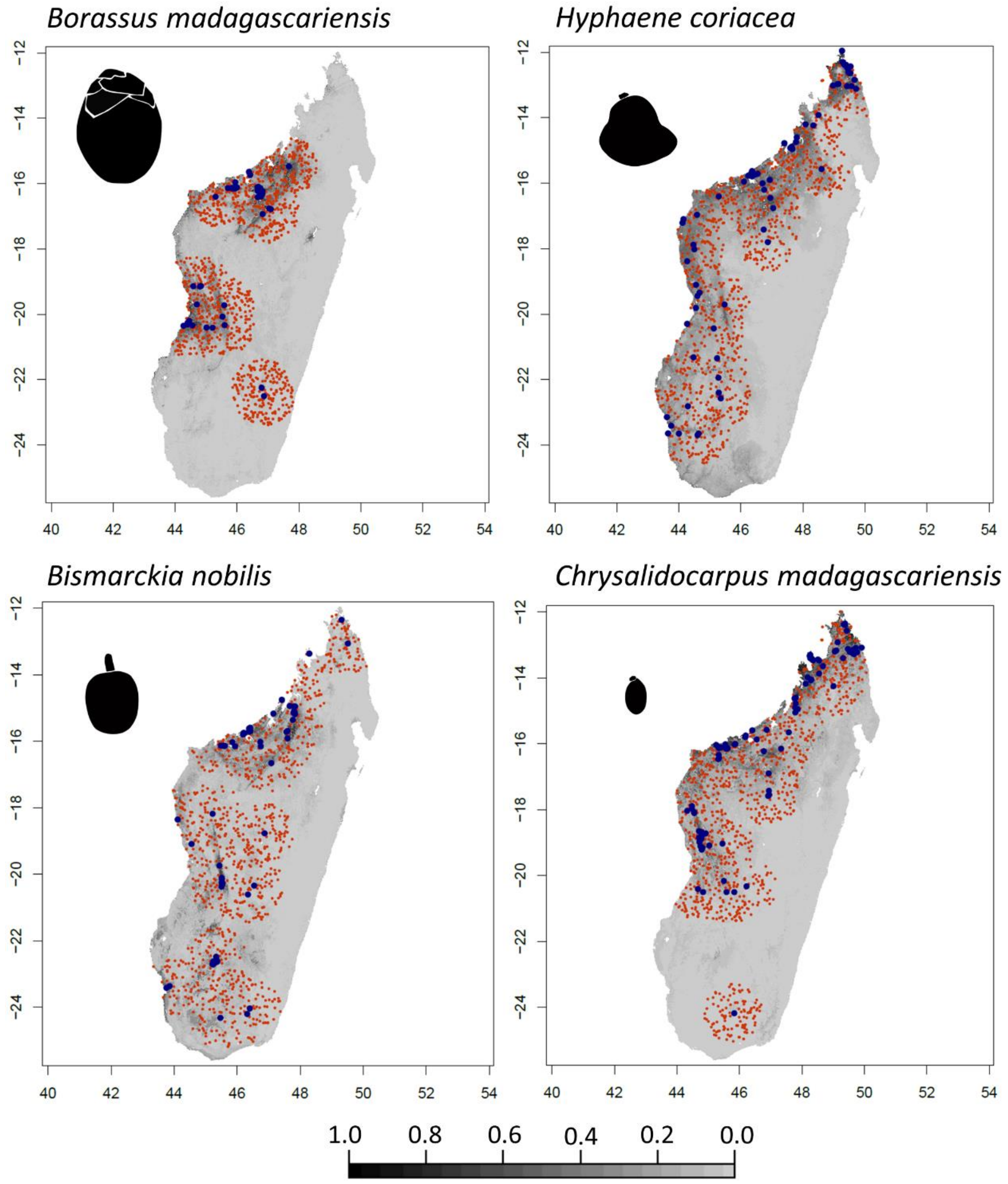


**Figure S2.** Occurrences and background points used to construct ensemble species distribution models for each palm species. Dark blue points represent thinned occurrences of each palm species (deleting points that are less than 3 Km from each other). Orange points represent background pseudo-absence points (1000 randomly generated points from within 100 Km of presence points).

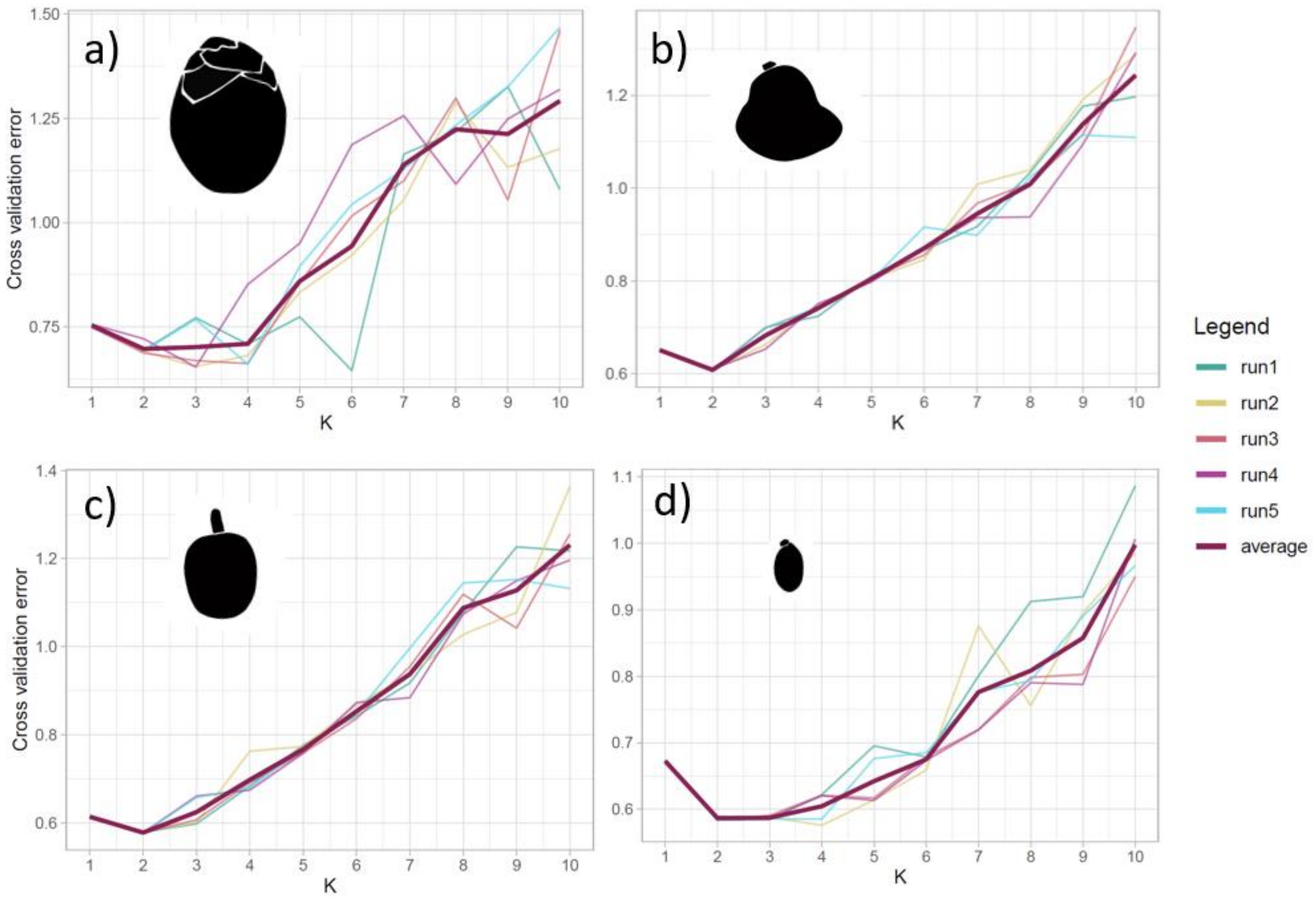


**Figure S3.** Cross validation error for the 5 different runs of ADMIXTURE and the average run (thicker, darker line) used for the choice of 'best' K for each species. a) *Borassus madagascariensis*, b) *Hyphaene coriacea*, c) *Bismarckia nobilis* and d) *Chrysalidocarpus madagascariensis*.

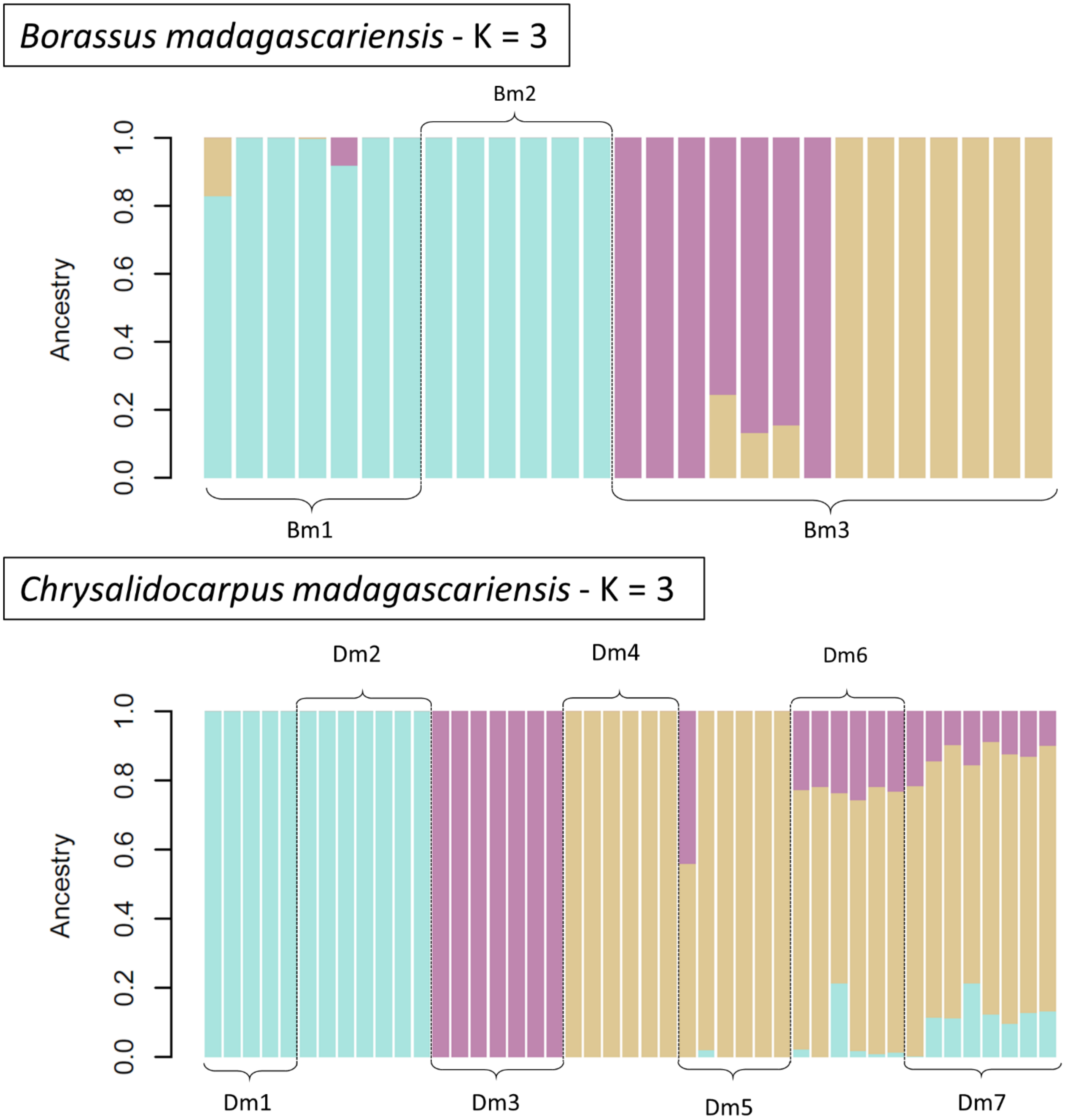


**Figure S4.** Results from ADMIXTURE analyses showing clustering K=3 for *B. madagascariensis* (Bm) and for *C. madagascariensis* (Cm)

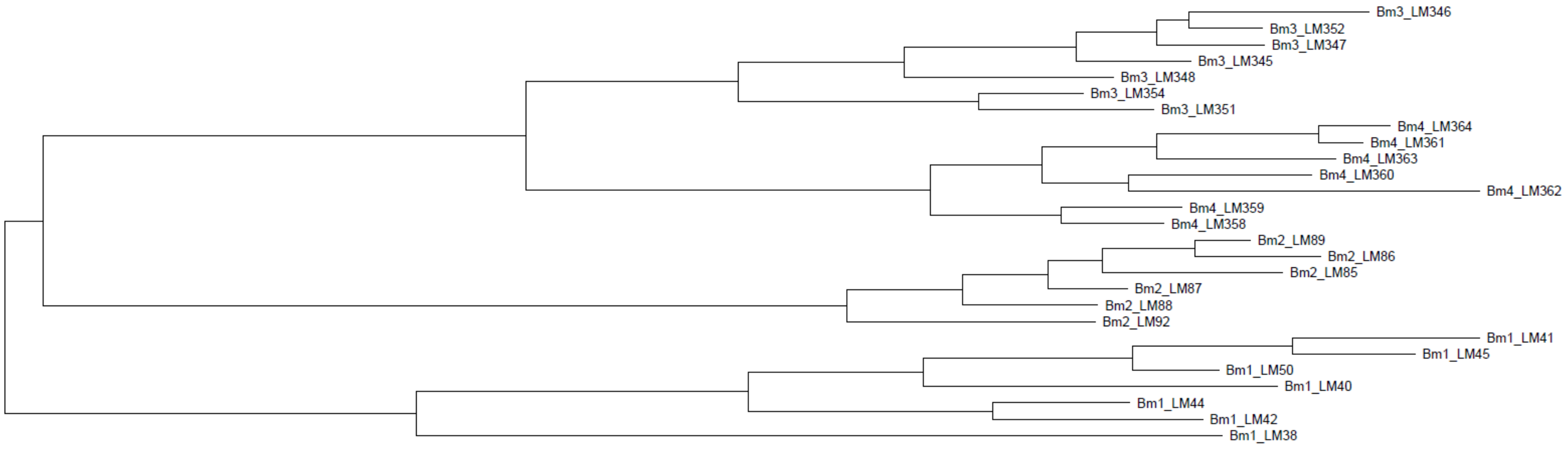


**Figure S5a.** Best tree for ***Borassus madagascariensis***. Tip labels correspond to all individuals sampled per population. In this case, individuals labeled as ‘Bm4’ were included into populations Bm3 since the individuals were sampled less than 2 km apart, but see Figure S4, where ADMIXTURE analyses also separated those two groups even though we considered them as the same population.

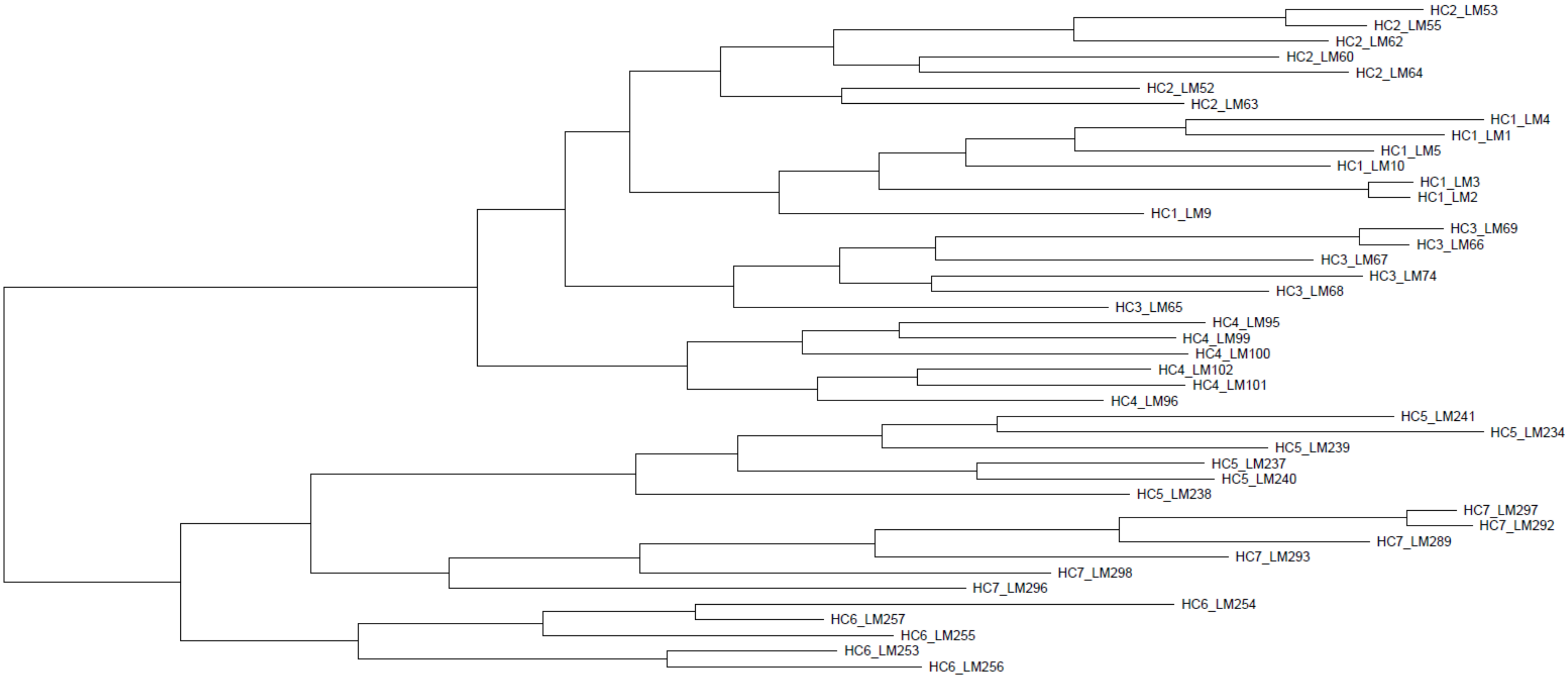


**Figure S5b.** Best tree for ***Hyphaene coriacea***. Tip labels correspond to all individuals sampled per population. All the natural populations cluster together.

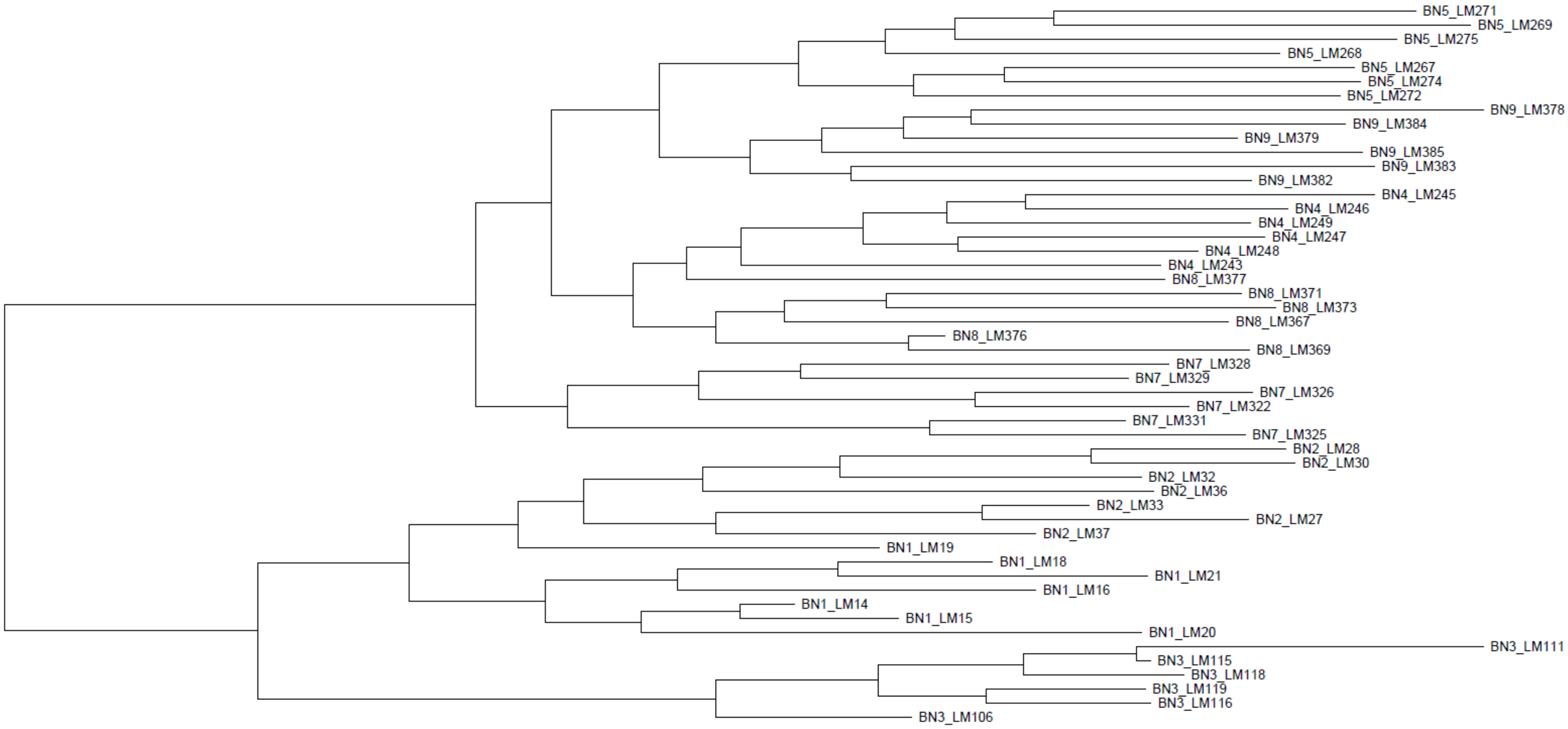


**Figure S5c.** Best tree for ***Bismarckia nobilis***. Tip labels correspond to all individuals sampled per population. Notice the admixture among populations BN1 - BN2 and BN4 - BN8.

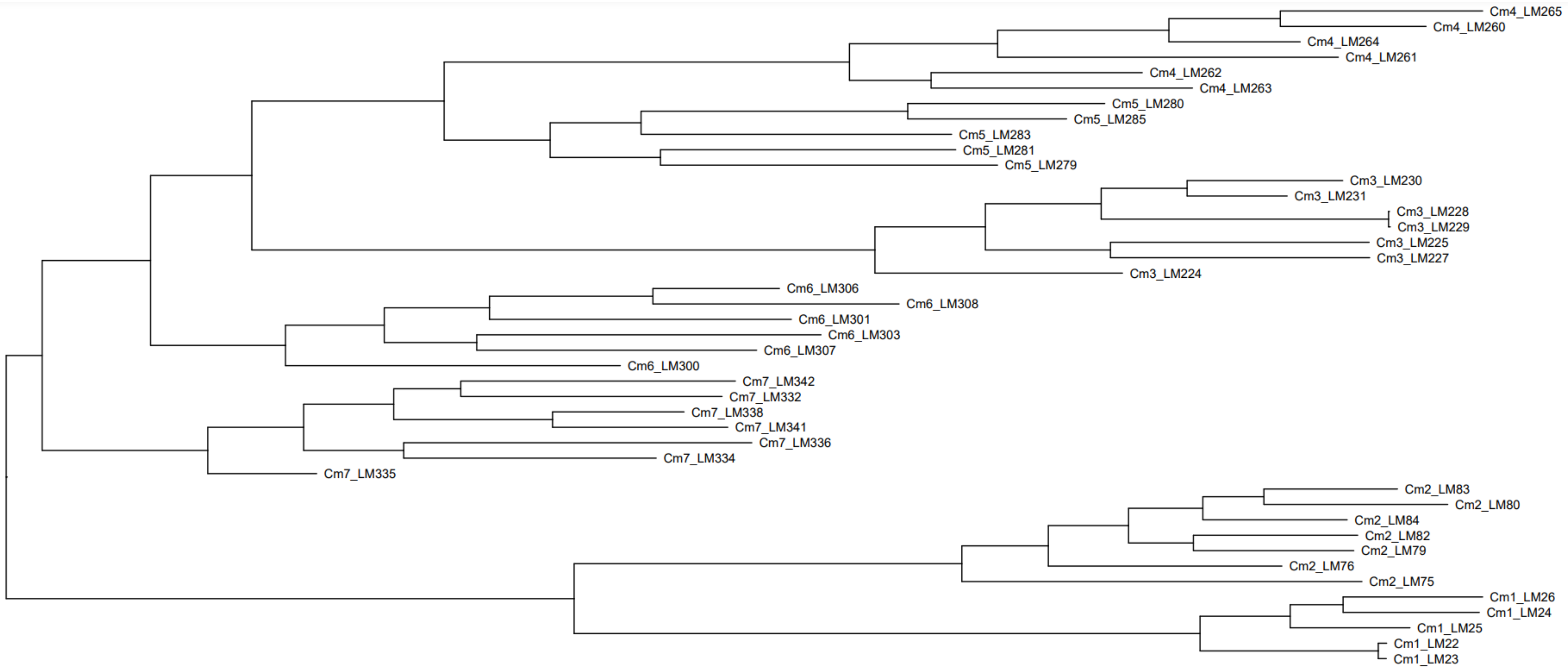


**Figure S5d.** Best tree for ***Chrysalidocarpus madagascariensis***. Tip labels correspond to all individuals sampled per population. All the natural populations cluster together.

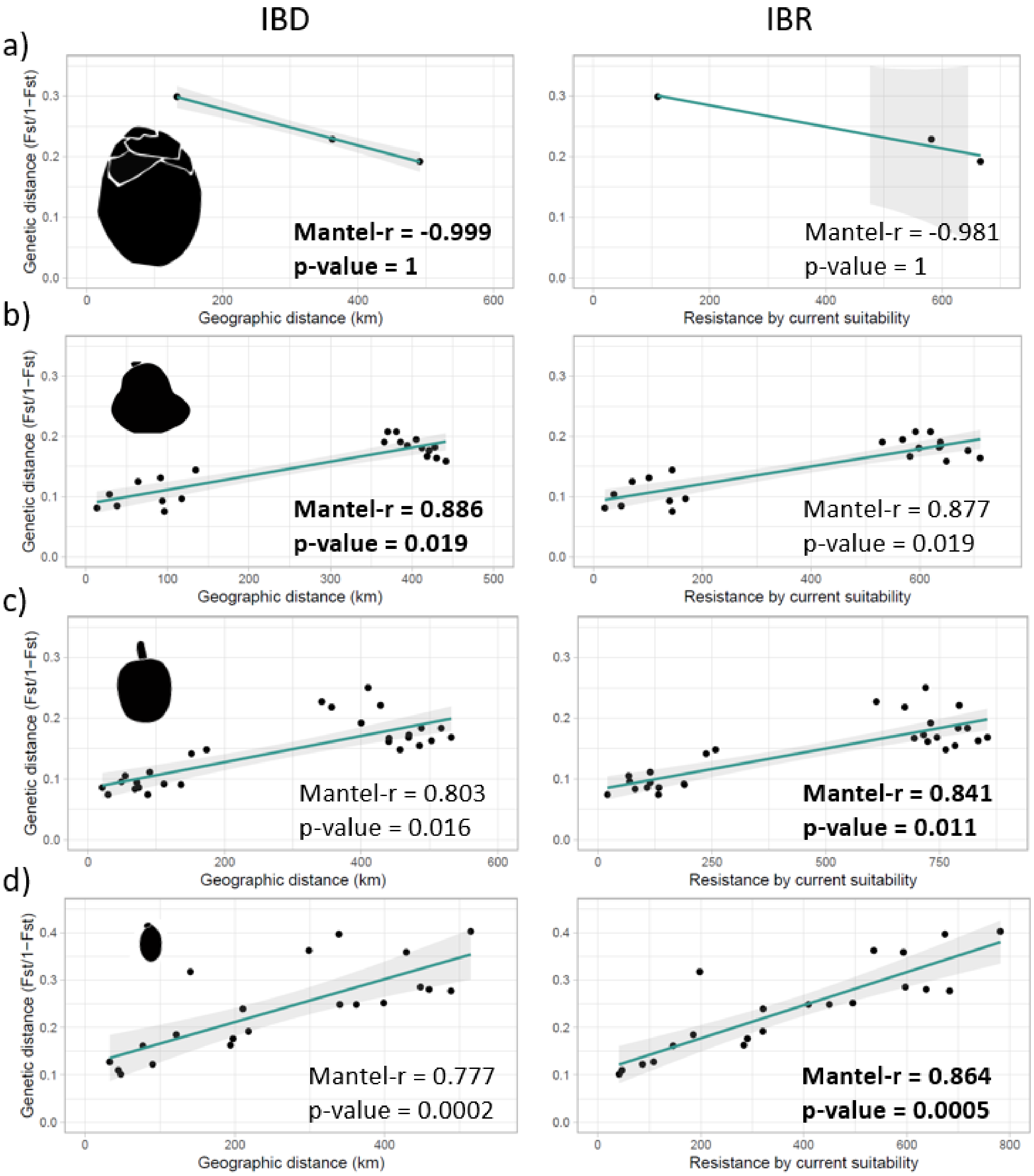


**Figure S6.** Results from the mantel tests of normalized genetic distance (Fst/1-Fst) or genetic differentiation between populations and isolation-by-distance (IBD) and isolation-by-resistance (IBR). Mantel-r or correlation is shown per species as well as p-values of each mantel test. The strongest correlations are shown in bold. a) *Borassus madagascariensis*, b) *Hyphaene coriacea*, c) *Bismarckia nobilis* and d) *Chrysalidocarpus madagascariensis*.

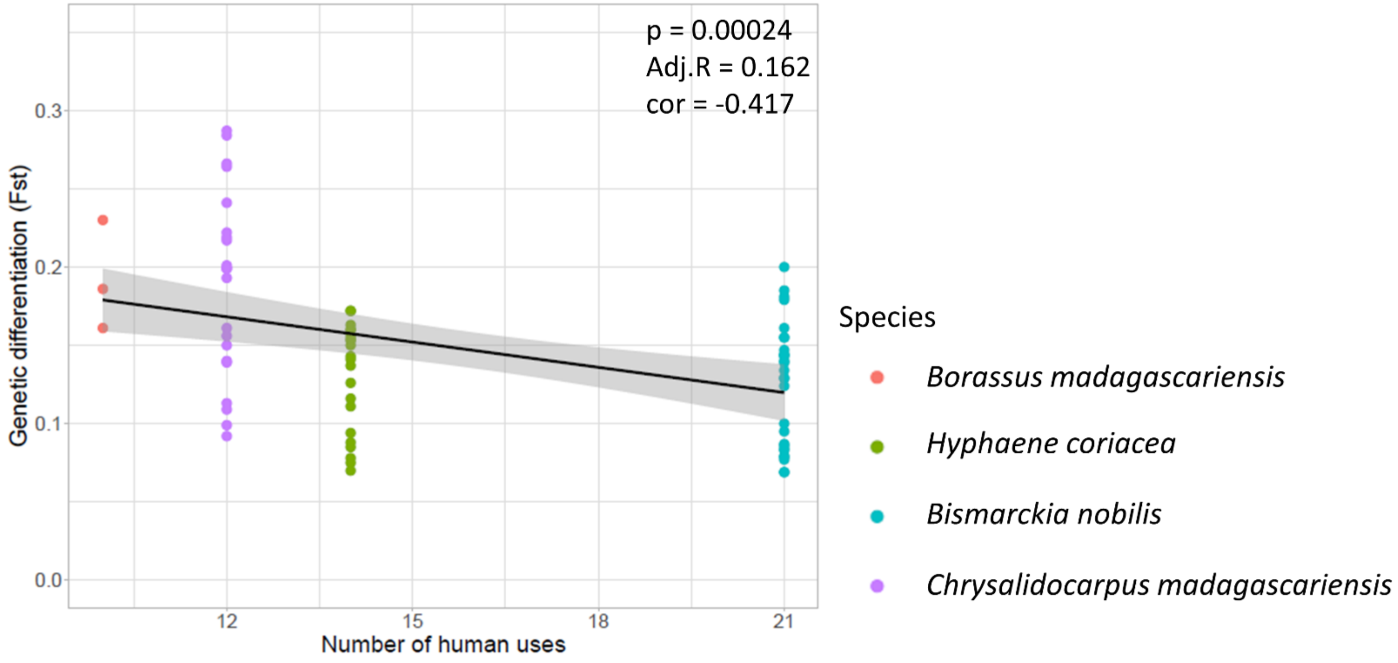


**Figure S7.** Correlation between genetic differentiation (Fst) of palms included in the study and number of uses by humans according to Rakotoarinivo et al. (2020). p = p-vale; Adj.R= adjusted R square; cor = correlation value.

### Supporting tables

**Table S1.** Coordinates of the centroids of all palm individuals of each population sampled in Madagascar and used in all subsequent analyses.

| Species | Population | Latitude | Longitude |
|---|---|---|---|
| *Borassus madagascariensis* | Bm1 | -20.3267381 | 44.4232619 |
| *Borassus madagascariensis* | Bm2 | -19.1424167 | 44.5874167 |
| *Borassus madagascariensis* | Bm3 | -16.1476111 | 45.9430417 |
| *Hyphaene coriacea* | HC1 | -19.7077083 | 45.4732361 |
| *Hyphaene coriacea* | HC2 | -19.4481389 | 44.5968611 |
| *Hyphaene coriacea* | HC3 | -19.3435556 | 44.6655278 |
| *Hyphaene coriacea* | HC4 | -19.1057222 | 44.5501667 |
| *Hyphaene coriacea* | HC5 | -16.7610000 | 47.0403333 |
| *Hyphaene coriacea* | HC6 | -16.0031389 | 46.6941389 |
| *Hyphaene coriacea* | HC7 | -15.9525833 | 46.0982778 |
| *Bismarckia nobilis* | BN1 | -20.1074167 | 45.5122222 |
| *Bismarckia nobilis* | BN2 | -20.3713492 | 45.5002381 |
| *Bismarckia nobilis* | BN3 | -19.0877222 | 44.5503333 |
| *Bismarckia nobilis* | BN4 | -16.1518944 | 46.7485833 |
| *Bismarckia nobilis* | BN5 | -15.6418056 | 46.3852500 |
| *Bismarckia nobilis* | BN7 | -16.1263611 | 45.4770000 |
| *Bismarckia nobilis* | BN8 | -16.1500000 | 45.9315278 |
| *Bismarckia nobilis* | BN9 | -15.7513611 | 46.2271111 |
| *Chrysalidocarpus madagascariensis* | Cm1 | -20.1613889 | 45.4994167 |
| *Chrysalidocarpus madagascariensis* | Cm2 | -19.1160873 | 44.7449524 |
| *Chrysalidocarpus madagascariensis* | Cm3 | -17.4250278 | 46.9418333 |
| *Chrysalidocarpus madagascariensis* | Cm4 | -15.5918333 | 46.4110278 |
| *Chrysalidocarpus madagascariensis* | Cm5 | -15.7959167 | 46.1865278 |
| *Chrysalidocarpus madagascariensis* | Cm6 | -16.0241111 | 45.8459722 |
| *Chrysalidocarpus madagascariensis* | Cm7 | -16.1164167 | 45.4115278 |

**Table S2**. Missing data per individual. Individuals in **bold** were excluded from subsequent analyses following the 'bad apples' rule (deleting individuals with more than 90% missing data).

| Individual | Number of variant sites | Number of missing sites | Percentage of missing data |
|---|---|---|---|
| | *Borassus madagascariensis* | | |
| Bm1_LM38 | 5091 | 1290 | 0.253388 |
| Bm1_LM40 | 5091 | 2016 | 0.395993 |
| Bm1_LM41 | 5091 | 1241 | 0.243764 |
| Bm1_LM42 | 5091 | 1340 | 0.26321 |
| Bm1_LM44 | 5091 | 1345 | 0.264192 |
| Bm1_LM45 | 5091 | 1291 | 0.253585 |
| Bm1_LM50 | 5091 | 1725 | 0.338833 |
| Bm2_LM85 | 5091 | 2429 | 0.477116 |
| Bm2_LM86 | 5091 | 2371 | 0.465724 |
| Bm2_LM87 | 5091 | 2368 | 0.465135 |
| Bm2_LM88 | 5091 | 2401 | 0.471617 |
| Bm2_LM89 | 5091 | 2406 | 0.472599 |
| Bm2_LM92 | 5091 | 2367 | 0.464938 |
| **Bm2_LM94** | **5091** | **4590** | **0.901591** |
| Bm3_LM345 | 5091 | 2840 | 0.557847 |
| Bm3_LM346 | 5091 | 2905 | 0.570615 |
| Bm3_LM347 | 5091 | 2842 | 0.55824 |
| Bm3_LM348 | 5091 | 2847 | 0.559222 |
| Bm3_LM351 | 5091 | 3319 | 0.651935 |
| Bm3_LM352 | 5091 | 2867 | 0.563151 |
| Bm3_LM354 | 5091 | 3404 | 0.668631 |
| Bm4_LM358 | 5091 | 2891 | 0.567865 |
| Bm4_LM359 | 5091 | 2869 | 0.563544 |
| Bm4_LM360 | 5091 | 4200 | 0.824985 |
| Bm4_LM361 | 5091 | 2841 | 0.558044 |
| Bm4_LM362 | 5091 | 4185 | 0.822039 |
| Bm4_LM363 | 5091 | 2893 | 0.568258 |
| Bm4_LM364 | 5091 | 2852 | 0.560204 |
| | *Hyphaene coriacea* | | |
| HC1_LM1 | 24503 | 9606 | 0.392034 |
| HC1_LM10 | 24503 | 9887 | 0.403502 |
| HC1_LM2 | 24503 | 9992 | 0.407787 |
| HC1_LM3 | 24503 | 10181 | 0.4155 |
| HC1_LM4 | 24503 | 9265 | 0.378117 |
| HC1_LM5 | 24503 | 9107 | 0.371669 |
| HC1_LM9 | 24503 | 9167 | 0.374117 |

| | | | |
|---|---|---|---|
| HC2_LM52 | 24503 | 11725 | 0.478513 |
| HC2_LM53 | 24503 | 12237 | 0.499408 |
| HC2_LM55 | 24503 | 12028 | 0.490879 |
| HC2_LM60 | 24503 | 11929 | 0.486838 |
| HC2_LM62 | 24503 | 11959 | 0.488063 |
| HC2_LM63 | 24503 | 12540 | 0.511774 |
| HC2_LM64 | 24503 | 13502 | 0.551035 |
| HC3_LM65 | 24503 | 15869 | 0.647635 |
| HC3_LM66 | 24503 | 15869 | 0.647635 |
| HC3_LM67 | 24503 | 15869 | 0.647635 |
| HC3_LM68 | 24503 | 15869 | 0.647635 |
| HC3_LM69 | 24503 | 15869 | 0.647635 |
| **HC3_LM73** | **24503** | **24503** | **1** |
| HC3_LM74 | 24503 | 15869 | 0.647635 |
| HC4_LM100 | 24503 | 14215 | 0.580133 |
| HC4_LM101 | 24503 | 14215 | 0.580133 |
| HC4_LM102 | 24503 | 14215 | 0.580133 |
| HC4_LM95 | 24503 | 14215 | 0.580133 |
| HC4_LM96 | 24503 | 14215 | 0.580133 |
| **HC4_LM98** | **24503** | **24503** | **1** |
| HC4_LM99 | 24503 | 14215 | 0.580133 |
| **HC5_LM233** | **24503** | **24503** | **1** |
| HC5_LM234 | 24503 | 12245 | 0.499735 |
| HC5_LM237 | 24503 | 12245 | 0.499735 |
| HC5_LM238 | 24503 | 12245 | 0.499735 |
| HC5_LM239 | 24503 | 12245 | 0.499735 |
| HC5_LM240 | 24503 | 12245 | 0.499735 |
| HC5_LM241 | 24503 | 12245 | 0.499735 |
| HC6_LM253 | 24503 | 6266 | 0.255724 |
| HC6_LM254 | 24503 | 9095 | 0.371179 |
| HC6_LM255 | 24503 | 5812 | 0.237195 |
| HC6_LM256 | 24503 | 6429 | 0.262376 |
| HC6_LM257 | 24503 | 5740 | 0.234257 |
| HC7_LM289 | 24503 | 14172 | 0.578378 |
| HC7_LM292 | 24503 | 14172 | 0.578378 |
| HC7_LM293 | 24503 | 14172 | 0.578378 |
| **HC7_LM294** | **24503** | **24502** | **0.999959** |
| HC7_LM296 | 24503 | 14172 | 0.578378 |
| HC7_LM297 | 24503 | 14172 | 0.578378 |
| HC7_LM298 | 24503 | 14172 | 0.578378 |
| *Bismarckia nobilis* | | | |

| | | | |
|---|---|---|---|
| BN1_LM14 | 21279 | 17395 | 0.817473 |
| BN1_LM15 | 21279 | 16630 | 0.781522 |
| BN1_LM16 | 21279 | 16644 | 0.78218 |
| BN1_LM18 | 21279 | 16642 | 0.782086 |
| BN1_LM19 | 21279 | 17346 | 0.81517 |
| BN1_LM20 | 21279 | 18020 | 0.846844 |
| BN1_LM21 | 21279 | 16816 | 0.790263 |
| BN2_LM27 | 21279 | 13786 | 0.647869 |
| BN2_LM28 | 21279 | 8789 | 0.413036 |
| BN2_LM30 | 21279 | 8845 | 0.415668 |
| BN2_LM32 | 21279 | 8696 | 0.408666 |
| BN2_LM33 | 21279 | 8672 | 0.407538 |
| BN2_LM36 | 21279 | 9459 | 0.444523 |
| BN2_LM37 | 21279 | 8758 | 0.411579 |
| BN3_LM106 | 21279 | 18173 | 0.854034 |
| **BN3_LM107** | **21279** | **21277** | **0.999906** |
| BN3_LM111 | 21279 | 18174 | 0.854081 |
| BN3_LM115 | 21279 | 18173 | 0.854034 |
| BN3_LM116 | 21279 | 18173 | 0.854034 |
| BN3_LM118 | 21279 | 18174 | 0.854081 |
| BN3_LM119 | 21279 | 18173 | 0.854034 |
| BN4_LM243 | 21279 | 15629 | 0.73448 |
| BN4_LM244 | 21279 | 16207 | 0.761643 |
| BN4_LM245 | 21279 | 15423 | 0.724799 |
| BN4_LM246 | 21279 | 15409 | 0.724141 |
| BN4_LM247 | 21279 | 15410 | 0.724188 |
| BN4_LM248 | 21279 | 16867 | 0.792659 |
| BN4_LM249 | 21279 | 16762 | 0.787725 |
| BN5_LM267 | 21279 | 9528 | 0.447765 |
| BN5_LM268 | 21279 | 14253 | 0.669815 |
| BN5_LM269 | 21279 | 9649 | 0.453452 |
| BN5_LM271 | 21279 | 9590 | 0.450679 |
| BN5_LM272 | 21279 | 9523 | 0.44753 |
| BN5_LM274 | 21279 | 9585 | 0.450444 |
| BN5_LM275 | 21279 | 10411 | 0.489262 |
| **BN6_LM310** | **21279** | **21279** | **1** |
| **BN6_LM313** | **21279** | **21279** | **1** |
| **BN6_LM315** | **21279** | **21279** | **1** |
| **BN6_LM316** | **21279** | **21279** | **1** |
| **BN6_LM317** | **21279** | **21279** | **1** |
| **BN6_LM318** | **21279** | **21279** | **1** |

| | | | |
|---|---|---|---|
| **BN6_LM319** | **21279** | **21279** | **1** |
| BN7_LM322 | 21279 | 17375 | 0.816533 |
| **BN7_LM323** | **21279** | **21278** | **0.999953** |
| BN7_LM325 | 21279 | 17375 | 0.816533 |
| BN7_LM326 | 21279 | 17375 | 0.816533 |
| BN7_LM328 | 21279 | 17375 | 0.816533 |
| BN7_LM329 | 21279 | 17375 | 0.816533 |
| BN7_LM331 | 21279 | 17375 | 0.816533 |
| BN8_LM367 | 21279 | 18968 | 0.891395 |
| BN8_LM369 | 21279 | 18968 | 0.891395 |
| BN8_LM371 | 21279 | 18969 | 0.891442 |
| BN8_LM373 | 21279 | 18968 | 0.891395 |
| **BN8_LM375** | **21279** | **21276** | **0.999859** |
| BN8_LM376 | 21279 | 18969 | 0.891442 |
| BN8_LM377 | 21279 | 18969 | 0.891442 |
| BN9_LM378 | 21279 | 12393 | 0.582405 |
| BN9_LM379 | 21279 | 12394 | 0.582452 |
| BN9_LM382 | 21279 | 12394 | 0.582452 |
| BN9_LM383 | 21279 | 12393 | 0.582405 |
| BN9_LM384 | 21279 | 12393 | 0.582405 |
| BN9_LM385 | 21279 | 12394 | 0.582452 |
| **BN9_LM386** | **21279** | **21275** | **0.999812** |
| *Chrysalidocarpus madagascariensis* | | | |
| Cm1_LM22 | 26847 | 6301 | 0.2347 |
| Cm1_LM23 | 26847 | 6356 | 0.236749 |
| Cm1_LM24 | 26847 | 7037 | 0.262115 |
| Cm1_LM25 | 26847 | 9319 | 0.347115 |
| Cm1_LM26 | 26847 | 7000 | 0.260737 |
| Cm2_LM75 | 26847 | 15988 | 0.595523 |
| Cm2_LM76 | 26847 | 19698 | 0.733713 |
| Cm2_LM79 | 26847 | 14909 | 0.555332 |
| Cm2_LM80 | 26847 | 14740 | 0.549037 |
| Cm2_LM82 | 26847 | 14744 | 0.549186 |
| Cm2_LM83 | 26847 | 14843 | 0.552874 |
| Cm2_LM84 | 26847 | 14730 | 0.548665 |
| Cm3_LM224 | 26847 | 11195 | 0.416993 |
| Cm3_LM225 | 26847 | 11079 | 0.412672 |
| Cm3_LM227 | 26847 | 11462 | 0.426938 |
| Cm3_LM228 | 26847 | 12211 | 0.454837 |
| Cm3_LM229 | 26847 | 11215 | 0.417738 |
| Cm3_LM230 | 26847 | 11012 | 0.410176 |

| | | | |
|---|---|---|---|
| Cm3_LM231 | 26847 | 13371 | 0.498044 |
| Cm4_LM260 | 26847 | 8435 | 0.314188 |
| Cm4_LM261 | 26847 | 8520 | 0.317354 |
| Cm4_LM262 | 26847 | 8421 | 0.313666 |
| Cm4_LM263 | 26847 | 14248 | 0.530711 |
| Cm4_LM264 | 26847 | 8369 | 0.311729 |
| Cm4_LM265 | 26847 | 9882 | 0.368086 |
| **Cm5_LM276** | **26847** | **26309** | **0.979961** |
| Cm5_LM277 | 26847 | 14423 | 0.537229 |
| Cm5_LM279 | 26847 | 14216 | 0.529519 |
| Cm5_LM280 | 26847 | 14239 | 0.530376 |
| Cm5_LM281 | 26847 | 14216 | 0.529519 |
| Cm5_LM283 | 26847 | 14217 | 0.529556 |
| Cm5_LM285 | 26847 | 14246 | 0.530637 |
| Cm6_LM300 | 26847 | 18280 | 0.680895 |
| Cm6_LM301 | 26847 | 19138 | 0.712854 |
| Cm6_LM302 | 26847 | 19581 | 0.729355 |
| Cm6_LM303 | 26847 | 18311 | 0.68205 |
| Cm6_LM306 | 26847 | 18249 | 0.679741 |
| Cm6_LM307 | 26847 | 18437 | 0.686743 |
| Cm6_LM308 | 26847 | 20466 | 0.76232 |
| Cm7_LM332 | 26847 | 16276 | 0.60625 |
| Cm7_LM334 | 26847 | 16275 | 0.606213 |
| Cm7_LM335 | 26847 | 17036 | 0.634559 |
| Cm7_LM336 | 26847 | 17396 | 0.647968 |
| Cm7_LM338 | 26847 | 19130 | 0.712556 |
| Cm7_LM341 | 26847 | 16261 | 0.605692 |
| Cm7_LM342 | 26847 | 16835 | 0.627072 |

**Table S3**. Information about number of individuals, populations sampled before and after deleting the 'bad apples'. Retained reads per species are shown. Number of single nucleotide polymorphisms (SNPs; only variant sites) recovered using all individuals or after deleting the 'bad apples', and before and after filtering with *populations* in Stacks v2.53 are provided. Notice that the number of SNPs recovered after deleting the 'bad apples' increases. Values in **bold** show the total number of SNPs per species that were used for all subsequent analyses (no bad apples, filtering with *populations* and only including sites with less than 50% missing data).

| Species | Number of populations | Number of individuals | Retained reads | Number of SNPs (variant sites) | Number of SNPs (variant sites) | Number of SNPs (variant sites) |
|---|---|---|---|---|---|---|
| **All individuals included** | | | | No filtering | Filtering with *populations* | |
| *Borassus madagascariensis* | 3 | 28 | 228464 | 183184 | 5091 | - |
| *Hyphaene coriacea* | 7 | 47 | 513399 | 649314 | 24503 | - |
| *Bismarckia nobilis* | 9 | 63 | 866044 | 1147571 | 21279 | - |
| *Chrysalidocarpus madagascariensis* | 7 | 46 | 665801 | 709425 | 26847 | - |
| **Only individuals with less than 90% missing data (no bad apples)** | | | | | Filtering with *populations* | Only sites with less than 50 % missing data |
| *Borassus madagascariensis* | 3 | 27 | - | - | 6324 | **1926** |
| *Hyphaene coriacea* | 7 | 43 | - | - | 28713 | **12898** |
| *Bismarckia nobilis* | 8 | 52 | - | - | 28728 | **5519** |
| *Chrysalidocarpus madagascariensis* | 7 | 45 | - | - | 28715 | **11589** |

**Table S4**. Model mean performance (five replicates per model) per species using presence only points and only the test dataset (30%). Numbers indicate Area Under the Curve of a Receiver Operating Characteristics plot (AUC) and True Skill Statistic (TSS) for each modelling algorithm. Values in **bold** represent models with mean AUC > 0.8 and mean TSS > 0.5, which indicates a good fit and therefore were used to construct the ensemble species distribution models.

| Modelling algorithm | Evaluation metric | |
|---|---|---|
| | AUC | TSS |
| *Borassus madagascariensis* | | |
| Boosted regression trees | **0.92** | **0.79** |
| Generalized additive models | 0.74 | 0.47 |
| Generalized linear models | **0.9** | **0.78** |
| Maximum entropy | **0.93** | **0.82** |
| Random forest | **0.92** | **0.8** |
| *Hyphaene coriacea* | | |
| Boosted regression trees | **0.82** | **0.53** |
| Generalized additive models | 0.78 | 0.5 |
| Generalized linear models | **0.83** | **0.57** |
| Maximum entropy | **0.84** | **0.57** |
| Random forest | **0.8** | **0.55** |
| *Bismarckia nobilis* | | |
| Boosted regression trees | **0.81** | **0.54** |
| Generalized additive models | **0.8** | **0.57** |
| Generalized linear models | 0.77 | 0.47 |
| Maximum entropy | 0.79 | 0.5 |
| Random forest | **0.82** | **0.57** |
| *Chrysalidocarpus madagascariensis* | | |
| Boosted regression trees | **0.87** | **0.64** |
| Generalized additive models | **0.84** | **0.59** |
| Generalized linear models | **0.86** | **0.58** |
| Maximum entropy | **0.89** | **0.66** |
| Random forest | **0.87** | **0.62** |

**Table S5**. Results from the Mantel tests between normalized genetic differentiation (Fst/(1 − Fst)) and distance between palm populations in km (isolation by distance – IBD) and least-cost distances between populations calculated from the current climate suitability maps resulting from the ensemble species distribution models (isolation by resistance – IBR). See also Table S8.

| Species | IBD | | IBR | |
|---|---|---|---|---|
| | **Mantel-r** | **p-value** | **Mantel-r** | **p-value** |
| *Borassus madagascariensis* | -0.999 | 1 | -0.981 | 1 |
| *Hyphaene coriacea* | 0.886 | 0.024 | 0.877 | 0.020 |
| *Bismarckia nobilis* | 0.803 | 0.013 | 0.841 | 0.009 |
| *Chrysalidocarpus madagascariensis* | 0.777 | 0.0001 | 0.864 | 0.0001 |

**Table S6.** ANOVA results using Tuckey Honest Significant Difference for genetic diversity (expected heterozygosity - He) and genetic differentiation (Fst) among the three fruit classes: large megafruits (*B. madagascariensis*), medium megafruits (*H. coriacea* and ***B.*** *nobilis*) and small fruits (*C. madagascariensis*). Difference = difference in the observed means; Lower = lower end point of the interval; Upper = upper end point of the interval; p adjusted = p-value after adjustment for multiple comparisons. In bold significant results are shown.

| Genetic diversity (He) | Difference | Lower | Upper | p adjusted |
|---|---|---|---|---|
| Large megafruits – medium megafruits | -0.0124 | -0.06103 | 0.036228 | 0.799518 |
| Large megafruits – small fruit | -0.04674 | -0.0998 | 0.00632 | 0.091113 |
| Medium megafruits – small fruit | -0.03434 | -0.06953 | 0.000858 | **0.05672** |

| Genetic differentiation (Fst) | Difference | Lower | Upper | p adjusted |
|---|---|---|---|---|
| Large megafruits – medium megafruits | -0.06578 | -0.12985 | -0.00171 | **0.042886** |
| Large megafruits – small fruit | -0.00419 | -0.07068 | 0.0623 | 0.987525 |
| Medium megafruits – small fruit | 0.061592 | 0.033494 | 0.089689 | **4.6E-06** |

**Table S7**. Genetic diversity per sampled population. Ho = Observed heterozygosity, He= Expected. heterozygosity, π= nucleotide diversity, Fis= inbreeding coefficient

| Population | Species | Private alleles | Number of individuals | Ho | He | π | Fis |
|---|---|---|---|---|---|---|---|
| Bm1 | *Borassus madagascariensis* | 184 | 5.96053 | 0.2489 | 0.24805 | 0.27146 | 0.0572 |
| Bm2 | *Borassus madagascariensis* | 210 | 4.94574 | 0.23332 | 0.19357 | 0.21614 | -0.03558 |
| Bm3 | *Borassus madagascariensis* | 362 | 10.45016 | 0.25469 | 0.25196 | 0.26473 | 0.02396 |
| BN1 | *Bismarckia nobilis* | 14 | 5.436 | 0.21861 | 0.20124 | 0.22205 | 0.00863 |
| BN2 | *Bismarckia nobilis* | 94 | 6.19538 | 0.21936 | 0.20892 | 0.22757 | 0.01798 |
| BN3 | *Bismarckia nobilis* | 53 | 4.63131 | 0.18401 | 0.15758 | 0.17729 | -0.01306 |
| BN4 | *Bismarckia nobilis* | 17 | 5.28989 | 0.23327 | 0.21487 | 0.23801 | 0.01061 |
| BN5 | *Bismarckia nobilis* | 40 | 5.48568 | 0.23475 | 0.22826 | 0.25171 | 0.03653 |
| BN7 | *Bismarckia nobilis* | 30 | 4.53706 | 0.23654 | 0.22089 | 0.24969 | 0.02866 |
| BN8 | *Bismarckia nobilis* | 21 | 4.59551 | 0.23593 | 0.21402 | 0.24102 | 0.01351 |
| BN9 | *Bismarckia nobilis* | 22 | 4.08855 | 0.22553 | 0.20753 | 0.24991 | 0.05058 |
| Cm1 | *Chrysalidocarpus madagascariensis* | 334 | 4.48877 | 0.13044 | 0.11332 | 0.12842 | -0.00371 |
| Cm2 | *Chrysalidocarpus madagascariensis* | 234 | 5.83359 | 0.12607 | 0.13182 | 0.14429 | 0.04113 |
| Cm3 | *Chrysalidocarpus madagascariensis* | 363 | 6.20697 | 0.19813 | 0.18529 | 0.20185 | 0.01028 |
| Cm4 | *Chrysalidocarpus madagascariensis* | 289 | 5.08393 | 0.22243 | 0.20097 | 0.22361 | 0.00354 |
| Cm5 | *Chrysalidocarpus madagascariensis* | 165 | 5.42711 | 0.24775 | 0.22781 | 0.25176 | 0.0092 |
| Cm6 | *Chrysalidocarpus madagascariensis* | 99 | 5.1561 | 0.24537 | 0.2226 | 0.24754 | 0.00471 |
| Cm7 | *Chrysalidocarpus madagascariensis* | 128 | 5.36964 | 0.23006 | 0.20938 | 0.23176 | 0.00508 |
| HC1 | *Hyphaene coriacea* | 54 | 6.48025 | 0.24206 | 0.23029 | 0.24977 | 0.01778 |
| HC2 | *Hyphaene coriacea* | 44 | 6.23692 | 0.24943 | 0.23714 | 0.25817 | 0.02079 |
| HC3 | *Hyphaene coriacea* | 42 | 5.05094 | 0.22687 | 0.22053 | 0.24556 | 0.03967 |
| HC4 | *Hyphaene coriacea* | 41 | 5.39727 | 0.2291 | 0.21718 | 0.23994 | 0.02313 |
| HC5 | *Hyphaene coriacea* | 246 | 4.63238 | 0.24155 | 0.2388 | 0.27132 | 0.06102 |
| HC6 | *Hyphaene coriacea* | 119 | 4.49592 | 0.27225 | 0.24592 | 0.27743 | 0.01054 |
| HC7 | *Hyphaene coriacea* | 163 | 4.72594 | 0.25026 | 0.23872 | 0.27182 | 0.04577 |

**Table S8**. Genetic differentiation (Fst) between pairs of populations, geographical distance between populations in kilometers and least cost paths between populations calculated from the current climate suitability maps resulting from the ensemble species distribution models. Bm = *Borassus madagascariensis*, HC = *Hyphaene coriacea*, BN = *Bismarckia nobilis* and Cm = *Chrysalidocarpus madagascariensis*.

| Population pairs | Genetic differentiation (Fst) | Geographic distance (Km) | Least cost paths/10000 |
|---|---|---|---|
| Bm1.Bm2 | 0.230 | 132.834 | 110.785 |
| Bm1.Bm3 | 0.161 | 490.978 | 666.150 |
| Bm2.Bm3 | 0.186 | 362.047 | 581.658 |
| HC1.HC2 | 0.070 | 96.254 | 144.616 |
| HC1.HC3 | 0.085 | 93.783 | 139.674 |
| HC1.HC4 | 0.088 | 117.555 | 168.882 |
| HC1.HC5 | 0.160 | 365.816 | 637.516 |
| HC1.HC6 | 0.141 | 429.909 | 710.716 |
| HC1.HC7 | 0.150 | 420.810 | 688.257 |
| HC2.HC3 | 0.075 | 13.640 | 20.635 |
| HC2.HC4 | 0.078 | 38.220 | 50.825 |
| HC2.HC5 | 0.156 | 394.114 | 638.461 |
| HC2.HC6 | 0.137 | 441.410 | 648.749 |
| HC2.HC7 | 0.143 | 418.369 | 581.616 |
| HC3.HC4 | 0.094 | 28.987 | 37.439 |
| HC3.HC5 | 0.172 | 380.666 | 619.384 |
| HC3.HC6 | 0.154 | 427.774 | 635.363 |
| HC3.HC7 | 0.163 | 404.915 | 568.230 |
| HC4.HC5 | 0.172 | 370.046 | 591.921 |
| HC4.HC6 | 0.153 | 411.952 | 597.924 |
| HC4.HC7 | 0.160 | 385.741 | 530.791 |
| HC5.HC6 | 0.116 | 91.659 | 101.724 |
| HC5.HC7 | 0.126 | 134.662 | 144.360 |
| HC6.HC7 | 0.111 | 64.029 | 71.089 |
| BN1.BN2 | 0.069 | 29.246 | 20.485 |
| BN1.BN3 | 0.124 | 151.409 | 237.300 |
| BN1.BN4 | 0.129 | 456.929 | 763.508 |
| BN1.BN5 | 0.140 | 502.831 | 834.647 |
| BN1.BN7 | 0.143 | 440.647 | 694.502 |
| BN1.BN8 | 0.139 | 440.256 | 723.853 |
| BN1.BN9 | 0.155 | 488.038 | 790.684 |
| BN2.BN3 | 0.129 | 173.512 | 257.785 |
| BN2.BN4 | 0.134 | 485.313 | 783.993 |
| BN2.BN5 | 0.144 | 531.788 | 855.132 |
| BN2.BN7 | 0.147 | 469.857 | 714.987 |

| | | | |
|---|---|---|---|
| BN2.BN8 | 0.144 | 469.454 | 744.338 |
| BN2.BN9 | 0.155 | 517.101 | 811.169 |
| BN3.BN4 | 0.161 | 399.987 | 730.343 |
| BN3.BN5 | 0.181 | 428.321 | 793.196 |
| BN3.BN7 | 0.185 | 342.187 | 610.944 |
| BN3.BN8 | 0.179 | 356.644 | 673.632 |
| BN3.BN9 | 0.200 | 409.964 | 719.197 |
| BN4.BN5 | 0.077 | 68.556 | 81.077 |
| BN4.BN7 | 0.083 | 136.038 | 189.616 |
| BN4.BN8 | 0.069 | 87.387 | 132.414 |
| BN4.BN9 | 0.086 | 71.283 | 114.280 |
| BN5.BN7 | 0.084 | 111.069 | 188.527 |
| BN5.BN8 | 0.079 | 74.319 | 133.379 |
| BN5.BN9 | 0.079 | 20.840 | 107.503 |
| BN7.BN8 | 0.087 | 48.687 | 68.963 |
| BN7.BN9 | 0.100 | 90.399 | 114.383 |
| BN8.BN9 | 0.095 | 54.289 | 67.063 |
| Cm1.Cm2 | 0.241 | 140.554 | 197.925 |
| Cm1.Cm3 | 0.284 | 338.906 | 674.048 |
| Cm1.Cm4 | 0.287 | 514.894 | 781.659 |
| Cm1.Cm5 | 0.217 | 488.620 | 683.129 |
| Cm1.Cm6 | 0.219 | 459.388 | 637.629 |
| Cm1.Cm7 | 0.222 | 447.804 | 597.120 |
| Cm2.Cm3 | 0.266 | 298.910 | 535.930 |
| Cm2.Cm4 | 0.264 | 428.966 | 593.550 |
| Cm2.Cm5 | 0.201 | 398.700 | 495.020 |
| Cm2.Cm6 | 0.199 | 362.226 | 449.521 |
| Cm2.Cm7 | 0.199 | 340.037 | 409.012 |
| Cm3.Cm4 | 0.193 | 210.634 | 320.794 |
| Cm3.Cm5 | 0.150 | 197.478 | 290.478 |
| Cm3.Cm6 | 0.140 | 194.143 | 283.517 |
| Cm3.Cm7 | 0.161 | 218.151 | 320.339 |
| Cm4.Cm5 | 0.113 | 33.002 | 108.349 |
| Cm4.Cm6 | 0.139 | 77.155 | 145.954 |
| Cm4.Cm7 | 0.156 | 121.783 | 185.257 |
| Cm5.Cm6 | 0.099 | 44.357 | 46.817 |
| Cm5.Cm7 | 0.109 | 90.232 | 86.727 |
| Cm6.Cm7 | 0.092 | 47.593 | 41.227 |

**Table S9.** Summary statistics after model-averaging used to explain genetic diversity (He, Ho, π) and genetic differentiation (Fst) of 25 populations of 4 species of palms throughout their distribution in the western part of Madagascar. Model averaged shows the conditional averaged estimates for the fixed effects and SE the standard errors. Sum of weights indicates the relative importance values of each explanatory variable, calculated over the best models which were selected by the model selection (ΔAICc ≤ 2). Number of containing models specifies in how many of the 'best models' after model selection each predictor occurred out of the total of best models. Percentage of forest was not significant explaining genetic diversity or genetic differentiation in any model. ***p ≈ 0; **p < 0.001; *p < 0.01.

| Predictor | **Genetic diversity** | | | | | | | | |
|---|---|---|---|---|---|---|---|---|---|
| | Expected heterozygosity (**He**) | | | Observed heterozygosity (**Ho**) | | | Nucleotide diversity (**π**) | | |
| | Model averaged (SE) | Sum of weights (%) | No. of containing models | Model averaged (SE) | Sum of weights (%) | No. of containing models | Model averaged (SE) | Sum of weights (%) | No. of containing models |
| Precipitation seasonality | 0.534 (0.153) *** | 100 | 3/3 | 0.496 (0.175) ** | 77 | 2/3 | 0.549 (0.150) *** | 100 | 4/4 |
| Road density | -0.276 (0.125) * | 30 | 1/3 | - | | | -0.282 (0.113) * | 27 | 1/4 |
| Temperature seasonality | -0.287 (0.150) . | 21 | 1/3 | -0.410 (0.183) * | 50 | 2/3 | -0.343 (0.131) * | 45 | 2/4 |
| Extant frugivore richness | - | | | - | | | -0.252 (0.115) * | 14 | 1/4 |

| Predictor | **Genetic differentiation (Fst)** | | |
|---|---|---|---|
| | Model averaged (SE) | Sum of weights (%) | No. of containing models |
| All four species | | | |
| Temperature seasonality | 0.469 (0.079) *** | 100 | 2/2 |
| Shared extinct megafrugivores | -0.258 (0.078) ** | 100 | 2/2 |
| Road density | 0.190 (0.068) ** | 68 | ½ |
| Only three large-fruited palm species | | | |
| Temperature seasonality | 0.605 (0.138) *** | 100 | 4/4 |
| Road density | 0.242 (0.081) ** | 84 | ¾ |
| Shared extinct megafrugivores | -0.311 (0.128) * | 68 | ¾ |
| Human population density | -0.254 (0.085) ** | 67 | 2/4 |
